# Statistical physics of liquid brains

**DOI:** 10.1101/478412

**Authors:** Jordi Piñero, Ricard Solé

**Affiliations:** ICREA-Complex Systems Lab, Universitat Pompeu Fabra, 08003 Barcelona, Spain; Institut de Biologia Evolutiva (CSIC-UPF), Psg Maritim Barceloneta, 37, 08003 Barcelona, Spain; Santa Fe Institute, 1399 Hyde Park Road, Santa Fe NM 87501, USA

**Keywords:** Brains, collective intelligence, criticality, phase transitions, evolution.

## Abstract

Liquid neural networks (or “liquid brains”) are a widespread class of cognitive living networks characterised by a common feature: the agents (ants or immune cells, for example) move in space. Thus, no fixed, long-term agent-agent connections are maintained, in contrast with standard neural systems. How is this class of systems capable of displaying cognitive abilities, from learning to decision-making? In this paper, the collective dynamics, memory and learning properties of liquid brains is explored under the perspective of statistical physics. Using a comparative approach, we review the generic properties of three large classes of systems, namely: standard neural networks (“solid brains”), ant colonies and the immune system. It is shown that, despite their intrinsic physical differences, these systems share key properties with standard neural systems in terms of formal descriptions, but strongly depart in other ways. On one hand, the attractors found in liquid brains are not always based on connection weights but instead on population abundances. However, some liquid systems use fluctuations in ways similar to those found in cortical networks, suggesting a relevant role of criticality as a way of rapidly reacting to external signals.

## I. INTRODUCTION

As pointed out by physicist John Hopfield, biology is different from physics in one fundamental way: biological systems perform computations, (Hopfield 1994). Within the context of evolution, a crucial ingredient for the emergence of biological complexity required the development of information-processing systems at multiple scales (Balušska & Levin 2015). Adaptation to a dynamic environment deeply benefited from non-genetic processes that allowed response mechanisms to short-term changes. Thus, biological computation is an intrinsic part of our current understanding of cell phenotypes (Benenson 2012) and not surprisingly the molecular webs of interactions connecting genes, proteins and metabolites has been often represented in terms of computations (Bray 1999).

Once fast-responding molecular signalling mechanisms were in place, a whole range of possibilities became available: individuals could not only respond to environmental cues, but they could also start to interact with other individuals prompting a higher-order cognitive network (Jablonka & Lamb 2006). Such transition took place in a diverse range of ways. It included the development of the first brain-like structures (Rose 2006, Pagan 2018) as well as societies formed by relatively simple agents (ants, termites or bees) capable of performing complex cognitive actions at the collective level (Oster & Wilson 1978; Solé & Goodwin 2001). Ant colonies have been compared to brains as both exhibit emergent collective phenomena (dynamical and structural patterns of organisation and behaviour that cannot be reduced to the properties of single ants) and display cognition on a large scale beyond that of the individual components (Gordon 1999, 2010). These two examples represent two distinguishable large classes of networks. Along with ant colonies, immune systems also share traits characteristic of the metazoan nerve nets yet they strongly depart from them in the fluid nature of cell-cell interactions.

The previous three examples are displayed in Figure 1. Here coupled neurons (a), interacting ants (b) or immune cells responding to novel challenges (c) are shown, along with minimal representations of the underlying networks (d-f). Here the classical picture of a neural network involves a topological structure (a graph) with neurons occupying the nodes and interneuronal links becoming the edges (d). Two types of nodes are shown, open and closed, associated to inactive or active neurons, respectively. Ant colonies, on the other hand, also involve collectives of interacting individuals which physical locations change over time: the colony is “liquid”. Thus, interactions are now limited to local neighboring agents, which constrains the system in a non-trivial manner. Within the liquid realm we can still characterize two paradigms: given their relative mobility and signal transmitivity (see below) inesct colonies are strongly affected by the locality of their interactions, whereas immune systems are highly mobile, such that a well-mixed approach might accurately represent their overall dynamics.

**Fig. 1:**
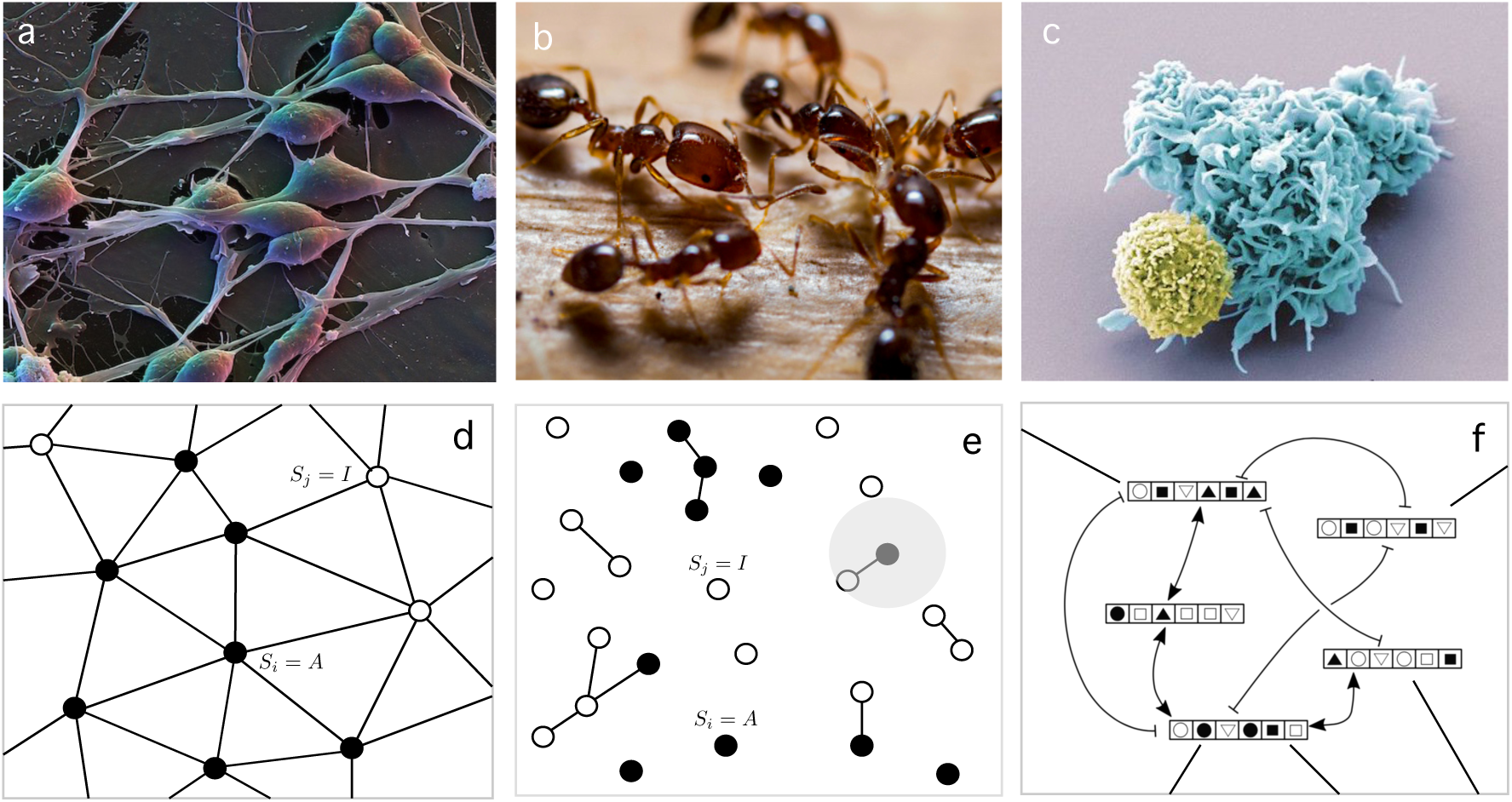
Network interactions in liquid versus solid brains. The three case studies analysed in this paper are shown, with examples of the agents involved in each case. Standard neural networks (a) involve spatially localized cells connected through synaptic weights. In contrast with this architecture, liquid brains, including (b) the immune system and (c) ant colonies include mobile agents (or cell subsets) interacting in space and time with no fixed pairwise weights. The schematic representation ofr each case study is outlined in the right column. Standard neural networks are defined in terms of connected excitable elements that can be roughly classified in active (firing) and inactive (quiescent) neurons, here indicated as filled and open circles, respectively (d). The wiring matrix remains basically the same in terms of topology (who is connected with whom) but will be modified in strength due to experience. By contrast, ant colonies must be represented by disconnected graphs (e) where interactions are possible within a given spatial range, here indicated by means of the grey circle. The immune system allows several representations of the interactions, but in many cases it is the molecular interaction between epitopes (strings of symbols in (f)) what truly represents the underlying liquid brain dynamics.

Other types of organisms, such as the slime mould *Physarum*, solve some classes of optimisation problems by utilizing a different form of fluid organisation (Tero *et al.* 2007) although in this case there is no neural substrate. This class of systems are able to solve minimization problems on a network (Adamatzky 2010).

Upon the transition to multicellularity, cell types capable of sensing and responding to signals appeared and permitted the emergence of a novel class of systems: webs of connected cells. These expanded the landscape of computations, including processing the information in nontrivial (aneural) ways (Balušska & Levin 2015). Such primitive networks provided a reliable way of dealing with complex decisions, integrating and storing memory and creating the conditions for increasing behavioral complexity. Simple organisms such as hydra and planarian flatworms provide good illustrations of the early steps in this direction (Pagan 2018). To some extent, all these systems can be modelled as networks of neurons that are connected in a stable way over time. Each pair of connected cells will remain linked over a given time scale and changes will take place at the level of the type and strength of the connection. Theoretical work has shown that cognitive tasks performed by these *solid* brains (simple and complex) such as pattern recognition, associative memory or language processing can be properly described. But what about liquid brains?

In this paper we review several models of both ant colony and immune system dynamics based on a neural network perspective and compare them with previous studies on “solid” brain models. In table I we summarise some general qualitative properties of the three classes of systems explored here, as well as others that we found relevant. The list is not exhaustive and involves generic descriptors that inevitably ignore the broad diversity of sizes, organization levels and ecological contexts. Several key components of each potential candidate, including size, age, context or developmental trajectories have some influence in the degree of robustness, memory potential or wiring patterns. All these factors make this basic table a tentative one. Nevertheless, it also highlights the commonalities that we consider relevant to our presentation.

Some key examples are worth mentioning. The label “liquid” is used to describe a physical state that ignores spatial structuring such as lymph nodes in the immune system or the nest structure of ant or termite colonies. Some of these features cannot be taken as absolute indicators since they are strongly influenced by life styles, size or behavioural context. The neural network of a hydra or a planarian flatworm are simple and small and might not display the modularity found in more complex neural agents, but nevertheless they display spatially stable networks of neurons, which are reliable under cell loss. Other relevant features (which are not included in Table 1) such as the self/nonself discrimination problem will be amply discussed later on. In the following sections, we summarise several types of models used to represent and understand the dynamics of the three case studies discussed here. By using them, we aim at enhancing the universal elements shared by these liquid systems while tracing a theoretical framework to study them.

**Table 1:** Comparative properties in Liquid *versus* Solid Brains. This table summarises a broad set of properties that are usually attributed to neural systems (soild brains) and here compared to those reported from two relevant examples of liquid brains, namely the immune system and insect (mostly ant) colonies. While the way computations are performed is a parallel process in all systems, all also exhibit some degree of specialisation, which can be understood as a modularity or a division of labour (DOL)- This first is observable in vertebrate brains while the later is a characteristic allocation of tasks that can occur either in societies with different morphological castes and in monomorphic ones. Similarly, we label the learning and memory properties in terms of a simple, network-related set of properties. In most cases studied here the memory potential of an ant colony is related to short-term phenomena tied to the production of a pheromone field, but long-term memories have also been reported at the individual level. In all these examples we indicate by (*) those attributes that are not well established or have been found in some case studies, and that will benefit from a theory of liquid brains.

|  | Brain | Immune System | Insect Colonies |
| --- | --- | --- | --- |
| Computational Nature | Distributed/Modular | Distributed/DOL | Distributed/DOL |
| Reliability under agent loss | High | High | High |
| Connectivity | Hard-wired yet plastic | Liquid (*) | Liquid but spatially constrained (*) |
| Memory & Learning | Synaptic | Population-based (Burnet) & Synaptic (Jerne) | Population-based & Synaptic |
| Regeneration potential | Low | High | High |
| Externalities | Peripheral Nervous System & Technology (tools) | No | Nests & Agriculture |
| Weighted Interactions | Hebbian | Antibody-Mediated | Pheromone-Mediated |
| Dynamical State | Critical | Critical (*) | Critical (*) |

## II. SOLID BRAINS

Standard neural networks (NN), from cell cultures to brains, have received great attention since the 1950s. A specially successful approach has been based on the use of statistical physics as a robust formalism capable of capturing the collective properties exhibited by neural masses (Deco *et al.* 2008). Both in statistical physics as well as in logic models of NN, neurons are replaced by a toy model representing only the minimal features exhibited by real cells. The intrincate structure of physiological neurons is ignored and replaced by a formal object devoid of any specific traits associated to cellular or molecular biological mechanisms. Similarly, the way connections and propagation of activity occurs is mapped into a simple graph. Despite all these oversimplifications, NN theory (also known as *connectionism*) has been capable of explaining the nature and relevance of collective phenomena involved in a broad range of areas, from learning in small metazoans to more complex phenomena related to human cognition (Forrest 1990, Farmer 1990).

We use here the term “brains” in a generic way too: it will refer to ensembles of interconnected neurons (or neural-like elements). The field has been growing since then into multiple directions, but a special turning point is the classical paper by Hopfield (Hopfield 1982) where the basis for a statistical physics description of neural networks emerged and largely marked the development of this class of systems. Such a “physics” perspective provided the basis for the understanding of their global properties out from the underlying microscopic description. Importantly, it also provided a systematic approach to identify the presence of different “phases” associated to the presence or lack of memory as well as dynamical states separating different types of activity. In this way, the physics of *phase transitions* (Amit *et al.* 1985, Sompolinsky 1988, Haken 1991) became a cornerstone to our understanding of neural networks.

The simplest, canonical model is based on an assembly of two-state agents description (McCulloch & Pitts 1943, Rashevsky 1960). These are denoted as *S*_*i*_(*t*) ∈ {0, 1} or *S*_*i*_(*t*) ∈ {−1, +1} (with *i* = 1, …, *N*). Agents are connected to each other through *fixed* synaptic links (Fig. 1a): each element sends to and recieves a signals from another. Connectivity is represented by a matrix *J*_*ik*_ ∈ *R*. The system is modelled by a dynamical set of equations:

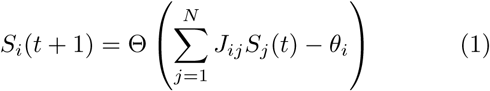

where Θ(*z*) = 1 for *z* > 0 and zero otherwise. The scalar *θ*_*i*_ is a threshold value. The so called *external field*, *h*_*i*_ = ∑_*j*_*J*_*ij*_*S*_*j*_(*t*) weights the total input of *S*_*i*_. It is worth noting that the same class of threshold model used to describe the dynamics of NN has been used to approach the dynamics of gene regulatory networks (GRN) (Kauffman 1993, Bornholdt 2005,2008).

### A. Attractor dynamics in recurrent neural networks

A general treatment of these systems involves a high-dimensional problem and a wide range of dynamical behaviours. However, an illustration of the potential of NN as a way of solving computational problems in a distributed manner is provided by the Hopfield model (Hopfield, 1982, Peretto 1992). This consists of a fully connected neural network described by the dynamical equations (1) with *θ*_*i*_ = 0. Hopfield’s model assumes no self-connection (*J*_*ii*_ = 0) and symmetry, i.e. *J*_*ij*_ = *J*_*ji*_. It can be shown that the model only displays single-point equilibrium (*attractors*), i.e., asymptotically, the trained network will tend to a stable configuration where all elements remain in a given state (Fig. 2a-c). Additionally, Hopfield’s model allows the network to store a number *p* of “memories” (patterns) defined as a set of vectors 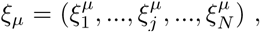, *µ* = 1 …, *p*. The storage process takes place within a “training phase” where they are presented to the network in such a way that each neuron *S*_*i*_ adopts the memory state i. e. *S*_*i*_ = *ξ*_*i*_ and all its synaptic weights *J*_*ij*_ are updated (starting from *J*_*ij*_ = 0 at time zero) following the so-called *Hebb’s rule*, which is summarised in Fig. 2b. In a nusthell, correlated inputs increase weights whereas uncorrelated ones decrease them. It can be shown (Hertz *et al.* 1991) that the memory states *ξ*_*µ*_ are, in fact, the minima of a (high-dimensional) energy function, namely:

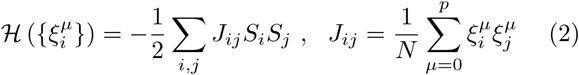

and initial conditions close to a minimum will evolve towards it. This is also outlined in Fig. 2c where we represent such multiple minima. In summary, the Hopfield model is a dynamical process of memory retrieval: stored patterns are recovered by a purely dynamical process. Extensions to this approach come by introducing thermal noise for the {*S*_*i*_} degrees of freedom. Usually, this is obtained via a *temperature T* that accounts for stochastic thermal variations (and, more generally, for noise). Each time we choose a neuron, the probability of changing to (or remaining in) state *S*_*i*_ = +1 is a saturating function

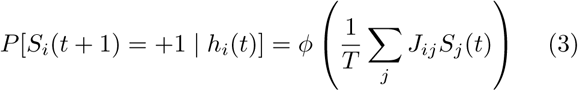

with *T* defining a temperature and *ϕ*(*x*) a function such that *ϕ*(0) = 0 and *ϕ*(*x*) → ±1 for *x* → ±∞. Temperature is not just an additional attribute, as it actually provides a powerful mechanism to escape from local minima. By using a stochastic transition rule, it is possible to move to lower-energy states from a given, suboptimal (usually non-memory) state. In this context, a measure of *memory capacity* is introduced as *α* ≡ *p/N*, where *p* here corresponds to the number of well-stored patterns. A phase-transition diagram captures the overall system behaviour, depicted in Fig. 1d. The shaded region represents states where the system is capable of retaining the memory patterns, while, for the blank region, these are lost due to noise. An abrupt transition separates these two regimes.

**Fig. 2:**
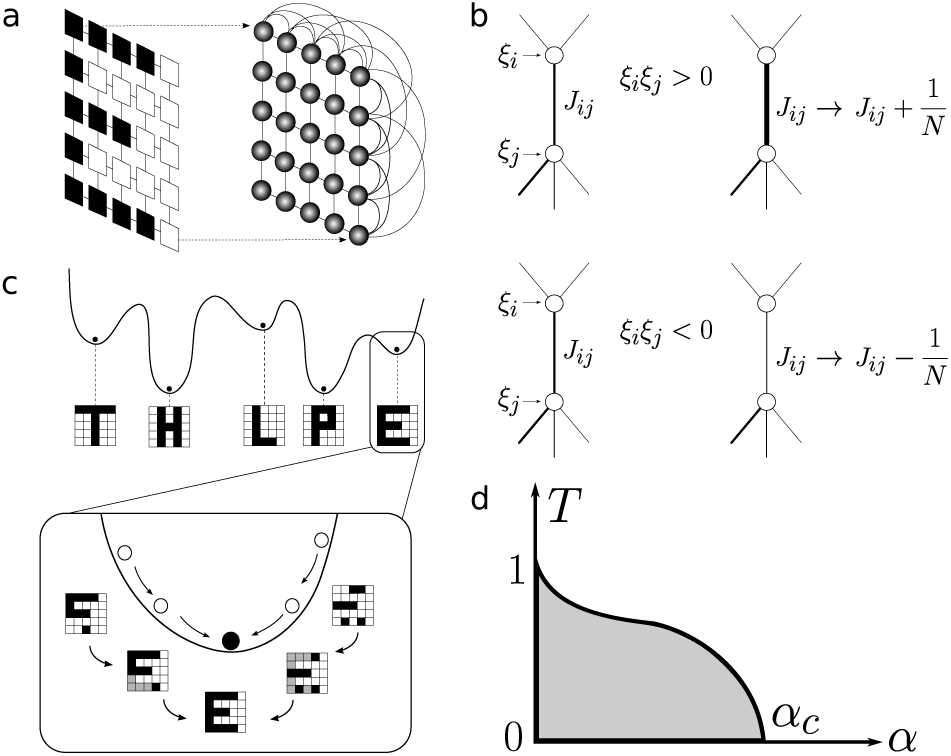
Distributed computation in neural networks. Using a very simple set of rules, a NN model can store and retrieve memories in a robust manner. In the Hopfield’s model, a massively connected set of neurons (a) with symmetric connections obeying Hebb’s rule (b) will display such properties. In (b), a pair of formal neurons is shown receiving inputs *ξ*_*i*_, *ξ*_*j*_ ∈ {−1, +1} from a given memory state or pattern *ξ*^*µ*^. If they are identical, i. e. *ξ*_*i*_ = *ξ*_*j*_, their connection is increased (in both directions). Otherwise, *J*_*ij*_ it is decreased. Network dynamics makes the system’s state flow to energy minima, thus recovering the desired memory state. The model exhibits remarkable reliability against connection loss. In (d) we show how reliable is memory retrieval against stochastic thermal variability. Parameter *α* is a relative measure of memory capacity. The critical value *α*_*c*_ ≃ 0.138 separates the two phases: memory reliability (shaded area) and unreliability (blank area). This transitions occurs sharply.

The previous model is an illustration of how cognitive functions can be understood in terms of a system of connected neurons. Here synaptic weights are modified in such a way that the resulting attractor dynamics allows associative memory to be the consequence of a relaxation towards energy minima. Only steady states are thus allowed. However, as discussed in the next section, a different picture emerges when we look at the actual dynamical patterns exhibited by neural tissues.

### B. Critical dynamics in cortical networks

If we think in an idealised graph such as the one described in Figure 1a, two classes of nodes can be defined: either inactive or active. Active nodes are formed by firing neurons whose excitability can be propagated to nearest inactive areas (Hesse & Gross 2014). As a result, excitation waves can move across whole areas. This would be a requirement to maintain integration in a dynamical fashion (Muñoz 2018). Instead of point, stable attractors are here replaced by more complex types of attractors.

The minimal model that can describe the propagation or activity is based on a contagion scenario where inactive nodes can become active if they are are connected to active nodes. Moreover, an active node can spontaneously decay. At the smallest scale, this is similar to the threshold dynamics described above. The simplest case to consider is a homogeneous model where all connections are similar, capable of propagating excitability with *J*_*ij*_ = *J* and an average connectivity 〈*k*〉. It can be shown that the large-scale (coarse-grained) dynamics for this homogeneous case can be defined by the equation (Hesse & Gross 2014):

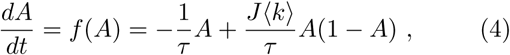

where *τ* is a characteristic time decay. A specially relevant observation is that neural systems exhibit *critical* behaviour (Chialvo 2004, 2010, Plenz *et al.* 2014). Two main classes of dynamical behaviour can occur. This can be shown using the fixed points, i.e., those *A*\* such that 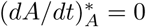. Two states are obtained. One is the trivial, inactive phase where no activity propagates: 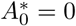. The second phase is associated to the second fixed point, namely:

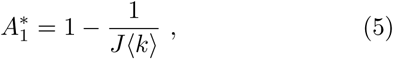

which is properly defined (i.e. 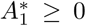) provided that *J*〈*k*〉 ≥ 1. A critical point separating the two *phases* is thus achieved for *J*〈*k*〉 = 1. For a given *J* value, the critical connectivity is given by 〈*k*〉_c_ = 1/*J*.

In Figure 3 two important diagrams are shown that summarise the basic phenomena resulting from the previous model. One is the so called bifurcation diagram (Strogatz 1994) where the stable states 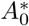, 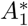 are plotted against the average connectivity *k*, with a marked change occurring at criticality. Additionally, we also display the potential function *V* (𝓐) (Solé 2011), defined as

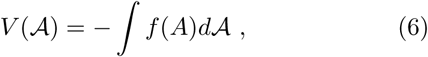

such that the dynamics derives from it, i.e. *d𝓐/dt* = −*dV* (𝓐)/*d𝓐*. The minima (maxima) of the potential correspond to stable (unstable) fixed points. As we approach criticality, the potential function becomes increasingly flatter. What is the impact of this flatness in the activity? In general, shallow potentials are associated to higher time variability and fluctuations diverge close to criticality. To show this, we can use a linear stability analysis taking the state 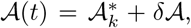, i.e. a small deviation *δ*𝓐 from a fixed point 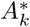, and pluggin it into the original equation for 𝓐(*t*). On a first approximation, it can be shown that

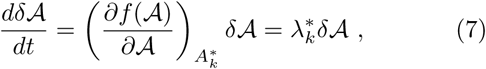

where 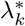 is a scalar to be evaluated at each fixed point. The resulting equation for fluctuations is linear. Thus, close to 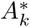, we expect a growth of fluctuations following an exponential growth or decay. For the inactive phase[99], we have

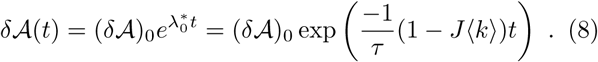

**Fig. 3:**
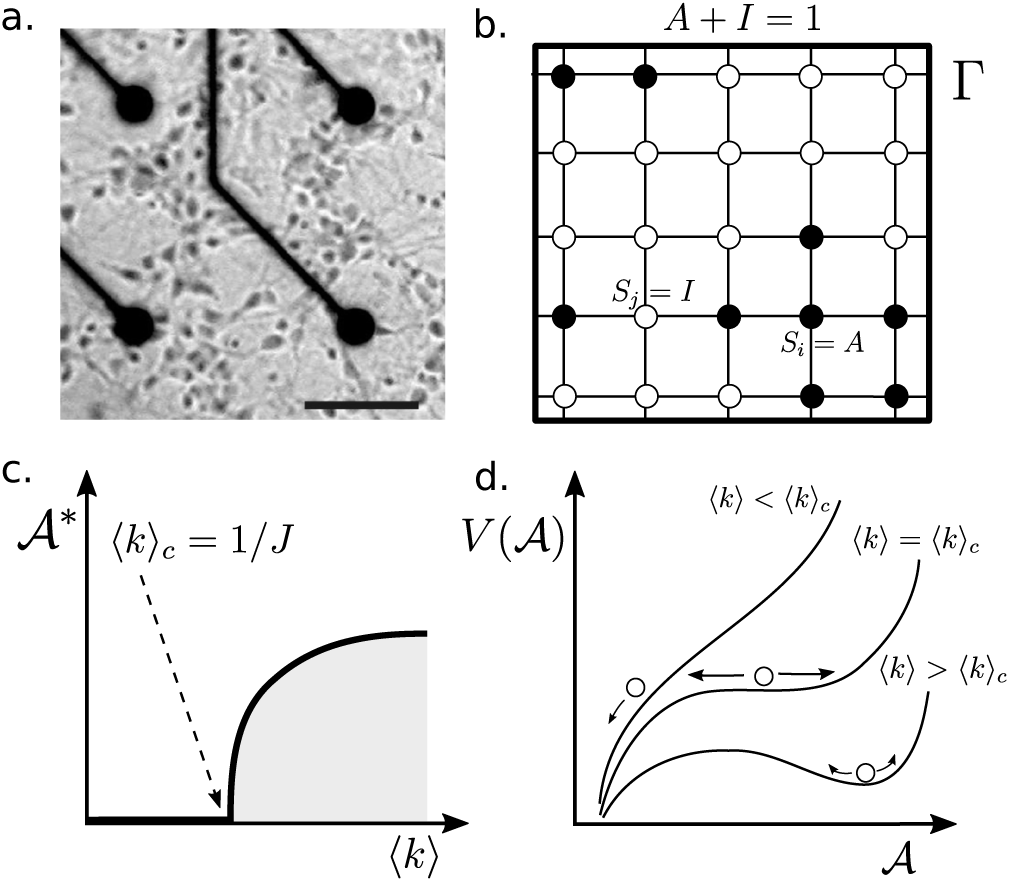
Phase transitions in neural dynamics. In a simple version of large scale dynamics of neural tissues (a) (such as brain cortex) can be represented as a network of connected neighbouring areas that are connected with excitatory links (adapted from Eckman *et al.* 2007). A toy model of this (b) could be represented as a lattice of neural elements connected as a grid with all elements linked to four elements in a homogenous fashion. Completar phase transitions. The analysis of this system (c-d) reveal a phase transition from zero activity to high-activity by crossing a critical value of average connections at 〈*k〉*_*c*_ = 1/*J*. A potential function can be obtained where the two phases are revealed as stable states of *V*(𝓐). Here, large fluctuations show clear dominance around the critical point.

As we can see the system will return to the fixed point (when 〈*k*〉 < 1/*J*) at a rate given by *λ*_0_. As we get close to criticality, the exponent gets smaller, the relaxation time rapidly increases. If the previous result is written in terms of a relaxation time *T* (*J*, 〈*k*〉), i. e. *δ*𝓐(*t*) *~* exp(*−t/T* (*J*, 〉*k*〈)) we have

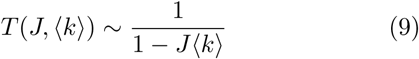

which rapidly diverges as *J*〈k〉 → 1. The divergence predicted by this simple model is confirmed by the analysis of the fluctuations found in neural systems.

The two previous models explore some essential components of neural complexity. Both deal with collective behaviour and exhibit special regions of parameter spaces that separate different phases. Phase transitions are of central importance within statistical physics, and provide a powerful framework to capture how microscopic interactions translate into system-level patterns and processes (Goldenfeld 1992, Solé 2011). Their importance within our context becomes manifest as qualitative changes in collective behaviour are typically caused by phase transition phenomena often associated to the density of individuals or the signals they use to communicate. How these systems behave close to transition points turns to be a key issue, as it provides understanding about how emergent phenomena occur.

## III. LIQUID BRAINS

### A. Ant colony dynamics

Social insects, including ants and termites among other groups, amount to about the same biomass than humans on Earth (Wilson 2012). With an evolutionary history spanning around a hundred million years, eusocial colonies have deeply engineered the environment and dominated the terrestrial biosphere much ahead (Wilson & Hölldobler 2005). In trying to attach biological fittness, insect colonies appear to behave as *superorganisms*, it is the colony as a whole that plays an evolutionary role, rather than its individual agents (ants). Across the biosphere, we encounter both monomorphic and polymorphic ant colonies. The latter involving physiological-anatomical differences within a given colony. However, it is estimated that 80% of ant species are monomorphic. The rest of species (polymorphic) range from 2 to 3 different casts. Here onwards, we will focus our study on monomorphic ant species.

On the other hand, various estimates state that the behavioral repertoire of ant colonies ranges from 20 to 45 different individual-ant behaviour (Oster & Wilson 1978; pp. 180-200). In order to shift from a given state to another and adapt to any given environmental circumstances, ants use chemical signals called *pheromones*. Different ant species use different sets of pheromones, some secrete only one type of molecule and others use up to twenty[100]. Thus, information is processed in a twolevel fashion: mobile agents (ants) interacting with a set of diffusive field of molecules (pheromones). Ants continuously detect the pheromone concentrations and, upon integrating this information, produce an internal image that affects their behavioural state. Moreover, ant states prompt the secretion of one (or more) pheromones thus reshaping their concentration values. This coalescence of signaling back and forth allows the whole colony to access global states where functions are achieved by means of its underlying network of interactions. Information is stored and processed through this “liquid brain” to give rise to various large scale collective behaviours. In the following examples, we will review several theoretical approaches to modelling ant colony dynamics and compare them with standard NN model efforts.

#### 1. Ant colonies as excitable neural nets

One of the simplest illustrations of the neural-like nature of insect colony dynamics is provided by the emergent synchronization displayed by some small colonies of the genus *Leptothorax*. In a nutshell, it has been observed that the colony-level activity displayed by their nests exhibits a remarkable bursting pattern (Fig. 4a,b) that exhibits a periodic component (Cole 1991). This means that ants can be active or inactive and the total number of active individuals changes in such a way that at times no ant in the colony is active while the synchronization events are linked to an almost fully active colony. These bursts have been found in other species (Hölldobler & Wilson 1990) result from the propagation of activity carried by moving individuals that can activate dormant ants in ways similar to those found in epidemic models (Goss *et al.* 1988, Bonabeau *et al.* 1998, Solé 2011). Synchronisation of neural masses is in fact a major research field within neuroscience (Buzsaki & Draguhn 2004) and it has been shown to pervade a wide range of functional traits and behavioural patterns. Is there something similar taking place in ant colonies?

**Fig. 4:**
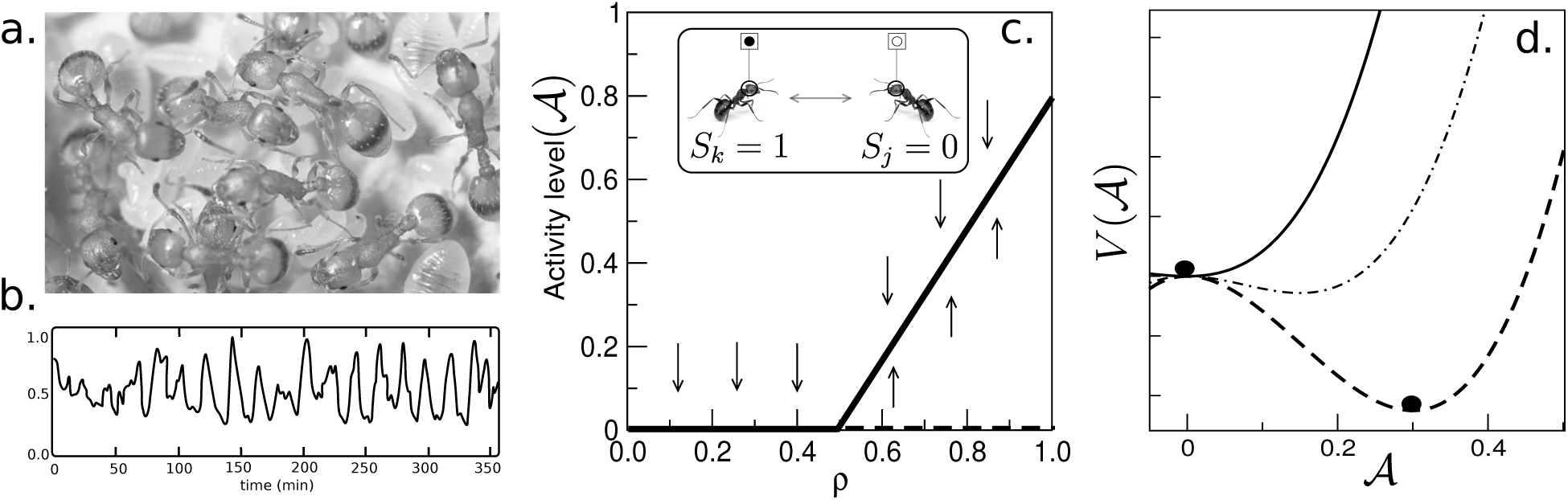
Ant colonies as excitable neural nets. In some ant species, such as those belonging to the genus *Leptothorax* (a), oscillations in activity have been recorded (b) revealing a collective synchronization phenomenon. This phenomenon can be described as an excitable neural system, where ants (inset of c) are reduced to a Boolean representation with active and inactive individuals, As the density of ants *ρ* increases, a phase change occurs (c) at a critical density, separating inactive from active colonies The potential function associated with the dynamics of these colonies is shown in (d): for densities larger (lower) than *ρ*_*c*_ is displays a well defined minimum. Closer to criticality, this potential becomes flatter and allows for wide fluctuations to occur.

This problem provides a simple example of a fluid network where the description level of individuals and their interactions is limited to a Boolean set of variables Σ = {0, 1} associated to the inactive (motionless) and active (moving) states, respectively (see inset of Fig. 4c). A NN model here is thus limited to a coarse-grained representation of ants. Such a model was suggested in (Solé *et al.* 1993) under the assumption that individuals can be described as an underlying continuous variable *S*_*i*_ ∈ [0, 1] (with *i* = 1, …, *N*) which change in time following a dynamical equation

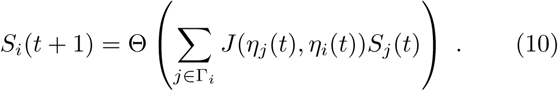

This dynamics strongly resembles the familiar form of standard NN. However a rapid inspection reveals a fundamental difference: here the matrix *J* (*η*_*j*_, *η*_*i*_) is state-dependent. In other words, its value is a function of the specific pair of agents that interact at a given time step. Specifically, we partition the activity interval [0, 1] into two domains associated to the active/inactive observables, i.e., *η*_i_ = Θ[*S*_*i*_ − *θ*]. Thus, the interaction matrix will include the only four possible pairs,

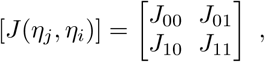

where *J* ≥ 0. Once activity decreases below the threshold *θ*, the ant becomes inactive and stops moving. Otherwise, it moves around as a random walker (unless constrained by other ants occupying nearest lattice sites). Here ants are assumed to move on a discrete two-dimensional lattice Ω and interactions occur in a strictly local manner, only affecting the set of nearest neighbouring positions Γ_*i*_ of *S*_*i*_. Finally, an inactive ant (with *η* < *θ*) can become active spontaneously (achieving a state *S*_0_ > *θ*) with probability *p*_*a*_. A common feature of these matrices is the presence of coupling terms connecting active and inactive individuals, as expected from an excitable system where activity can be propagated among agents. It is important to notice that the collective synchronisation does *not* result from the coupling of individuals’ internal clock. Instead, single virtual ants behave randomly. The dynamics of single elements will be described by:

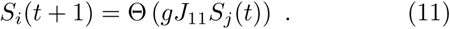

A simple case can be solved, namely when the coupling is small and activity remains small (which is consistent with observation). If we choose Θ(*x*) = tanh *x*, then we may use linear approximation tanh(*gJz*) ≈ *gJz* which admits a solution to the previous equation. If, initially, an ant is activated to a level *S*_0_, then *S*(*t*) = *S*_0_(*gJ*)^*t*^, which is a decaying function of time. If an activation term is also introduced (i.e. active ants can activate inactive ones), then a coarse grained model can be defined in probabilistic terms. Let us label as *N*_*a*_ the number of active ants. This number will change in time as a consequence of both interactions and decay. The efficiency of activation events will be proportional to *gJ*, assuming the previous linear approximation. Hereafter we will indicate by *N* and *ρ* the total number and density of ants, respectively.

If 𝓐(**x**, *t*) indicates the probability density of active ants at a given point of our two-dimensional lattice **x** ∈ Ω, then it can be written as: 𝓐(**x**, *t*) = *P* [*S*_**x**_(*t*) = 1]. The activity density will evolve following a master equation according to the previous rules:

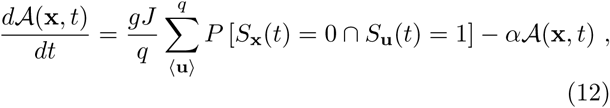

where 〈**u**〉 indicates sum over the set of *q* nearest neighbors, *P* [*S*_**x**_ = 0 ∩ *S*_**u**_ = 1] is the probability of having a pair o nearest ants in different states.

The previous equation is exact, but its computation would require knowledge of the probabilities associated with the interactions between nearest sites. Several methods can be used to solve this model with different levels of approximation. Here we will consider the simplest one, commonly known as a *mean field theory*, which is based on suppressing the spatial correlation between nearest sites. This is done by assuming that the system is in fact well mixed and thus all sites are neighbours or, in mathematical terms, *q* = Ω. If this is the case, we can use the total population

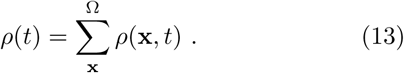

By summing on both sides of the previous master equation, and ignoring correlations between active and inactive neighbours, it can be shown that the global dynamics can be described as:

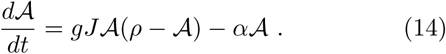

And this equation can be studied as a deterministic model of ant colonies displaying excitable dynamics. The model has two equilibrium points, namely 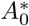 = 0 (no activity spreads) and 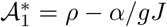, associated to persistent propagation. The previous equation is similar to those used in epidemic dynamics (Murray 1989) associated to a population composed by two classes of individuals (infected and susceptible). Using the density of ants as a control parameter, these two phases are separated by a critical point *ρ_c_* = *α/gJ*. The global behaviour of this model is summarised in Figure 4c where the bifurcation diagram for this system is shown. Above *ρ*_*c*_ an active phase is present whereas an inactive one if found for *ρ* < *ρ*_*c*_.

In this system, the potential function *V* (𝓐) is:

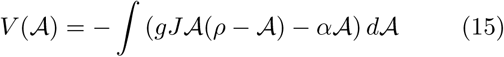

and is displayed in Figure 4d, where we show three examples of its behaviour for different density values. As we already discussed within the context of brain criticality, here too the transition between phases as density is changed involves a shallow potential function, indicating thet wide fluctuations should be expected to occur. One remarkable observation from *Leptothorax* colonies is that they seem to be poised close to the critical density (Miramontes 1995) at density levels where theory predicts that maximum information and behavioural diversity is achieved (Solé & Miramontes 1995, Miramontes & DeSouza 1996). As discussed above within the context of neural tissues, criticality provides a source of fast response and optimal information processing.

The key message provided by this example is that a commonality with other excitable neural systems exists: a universal property is the use of critical points to perform cognitive tasks. Being poised close to critical states provides a natural way of amplifying input signals while remaining most of the time in a low-fluctuation state (Mora & Bialek 2011). Such a compromise makes sense as a way of displaying optimal information while reducing the cost of the system’s state. Is there a well-defined function that can be associated to this? The answer is yes. By using self-synchronized patterns of activity a task may be fulfilled moreeffectively than with non-synchronised activity, at the same average level of activity per individual Delgado & Solé 1997a, 2000).

#### 2. Collective decision making and symmetry breaking in ant colonies

The next case study involves one of the best examples of how fluid brains solve a well-defined optimisation problem. Specifically, a given ant colony exploiting a number of sources of nutrients might need to discriminate between different sources (Deneubourg & Goss 1989, Detrain & Deneubourg 2006, Garnier *et al.* 2007). Another problem (which we explore here) involves the determination of the shortest path to be chosen between two alternatives. This problem can be easily implemented in the lab, using a two-bridge setup (Figure 5a). Here the ant nest would be located in the left side and ants would walk through the two-bridge to reach a food source located on the right side. The two branches can be identical or instead have different lengths. The problem to be solved here is which one is the shortest. Once again, the solution cannot be found at the individual level: colony-level processes need to be in place to make the right decision.

**Fig. 5:**
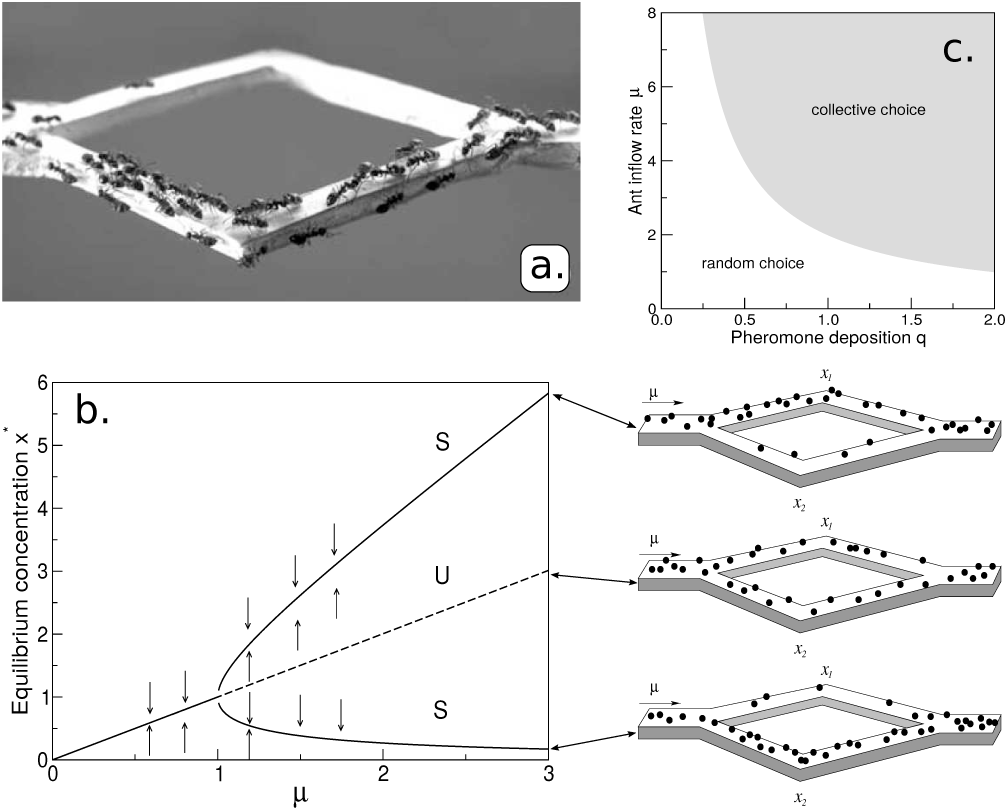
Collective decision making. A two-path experiment (a) allows to test the mechanisms by which emergent decision making occurs. The photograph shows an example of a colony that has made a collective decision, as shown by the preferential use of the shortest one. (b) The mathematical analysis of the model associated to this phenomenon shows that two alternative solutions exist associated to the preferential choice of one branch, along with a third one where both branches are used. In (c) the parameter space for the simple symmetric case is shown.

Ants can use quorum-sensing mechanisms as a way of creating and (responding to) pheromone fields thus generating a large-scale chemical field that allows to properly perform the decision. Initially, ants will walk on both bridges, choosing randomly their branch. We should expect at this point equal number of ants on each branch, i.e. *ρ*_1_ = *ρ*_2_. However, once an ant has found the food source, it releases a pheromone as it returns to the nest. Other ants will detect the released signal, which helps ants to decide where to move, releasing further pheromones and amplifying the previous mark. The pheromone trail also evaporates, and evaporation will be more effective in the longer trail, where more surface is available. As a result, the shortest path is more likely to be used, and is eventually chosen. Ants have computed the shortest path. A model describing this experiment can be defined as follows. If *ρ*_1_ and *ρ*_2_ indicate the concentrations of trail pheromone in each branch, their dynamics (Nicolis & Denebourg 1999) is given by a pair of equations for the pheromone fields:

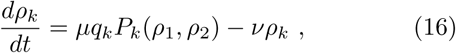

with *k* = 1, 2. Here *µ* is the rate of ants entering each branch, *q*_*i*_ the rate of pheromone production at the *i*-th branch and *ν* is the rate of evaporation. The functions *P*_*i*_(*ρ*_1_, *ρ*_2_) can now be understood as probabilities of choosing a bridge depending on thepheromone concentrations. These probabilities are well described by a nonlinear, threshold response function (Beckers *et al.* 1992, Deneubourg *et al.* 1990):

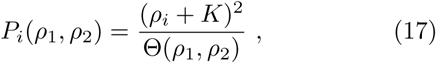

where Θ(*ρ*_1_, *ρ*_2_) = ∑_*j*=1,2_(*ρ*_*j*_ + *K*)^2^ and *i* = 1, 2. The parameter *K* gives the likelihood of choosing a path free of pheromones (*ρ*_i_ = 0).

This is a general model that incorporates attributes associated to each branch. But an interesting scenario arises when one considers the symmetric case where *q*_1_ = *q*_2_ = *q*. For this situation the previous set of equations

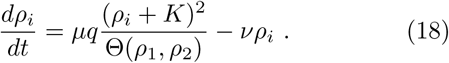

Here there is no true optimal choice: both branches are equal. Now, although the obvious expectation is a similar disitribution of ants in each branch, this is not what is observed. We would easily conclude that ants would choose both paths and that individuals will equally walk in both branches. However, what is typically seen is that the symmetry is broken in favour of one of the two branches. Why is this the case? This phenomenon illustrates a very important class of phase transition: the so called *symmetry breaking* process. Despite the symmetry of the system, amplification of initial fluctuations leads to the formation of a dominant pheromone trail that is used by all ants once established.

The fixed points associated to this system are obtained from *dρ*_*i*_/*dt* = 0. One possible solution to this systemis the symmetric state 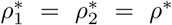 (associated to equal use of both branches)ants equally distributed in both branches). For this special case, we have 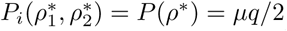 and thus a we only need to solve a single equation *dρ*^*^/*dt* = *µq*/2 − *νρ*^*^, which gives a fixed point *ρ*^*^ = *µq*/(2*ν*). This is the symmetric state to be broken. The second scenario corresponds to the choice of one of the branches 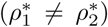 *ρ*_1_ + *ρ*_2_ = 2*ρ*\* = *μq/ν*, we see that

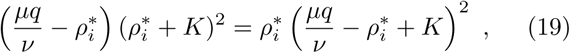

after some algebra, this gives the new fixed points 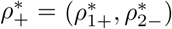 and 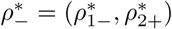 with

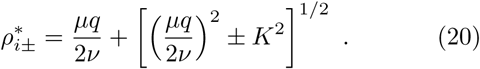

This pair of fixed points will exist provided that *µq*/2*ν* > *K* which allows to derive a critical line (Fig. 5c)

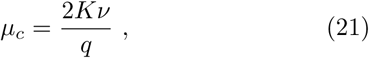

indicating that there is a minimal rate of ants entering the bridges required to observe the symmetry breaking phenomena. For *µ* > *µ*_*c*_ the symmetric state becomes unstable (see Figure 5b-c) while the two other solutions can be equally likely. Below this value, the only fixed point is the symmetric case withidentical flows of ants in each branch. This symmetric model can be generalized to (more interesting)asymmetric scenarios where the two potential choices are different (see Detrain & Deneubourg 2006 and references therein) either because the food sources have different size or because paths have different lengths and the shortest path need to be chosen. This symmetry breaking phenomenon has also been observed in the ant colony panic responses (Altshuler et al 2005) or army ant trails (Deneubourg et al 1989) or optimal group formation (Amé et al 2006).A specially interesting proposal concerning the phenomenon of symmetry breaking in ants was made in (Bonabeau 1996), where it was suggested that flexible behaviour leading to efficient decisions is more likely to occur close to critical points.

#### 3. Task allocation in ant colonies as a parallel distributed process

In the previous example we considered a set of agents described as binary variables, thus ignoring the combinatorial complexity that should be expected from an insect equipped with a brain. Moreover, it is clear that the active/inactive dichotomy hides a repertoire of potential activities that can be carried out by individuals, associated to the set of tasks needed to maintain the colony. Division of labour is in fact one of the most important and widespread phenomenon in nature, and very common in social groups (Duarte *et al.* 2011). It has been shown that the dynamics of subsets of individuals performing specific tasks within colonies is an emergent phenomenon (Gordon 1999). In this scenario, a colony that needs to perform a given set of tasks under given environmental conditions (and respond to changes in flexible ways) must be capable of sensing its internal state using some kind of distributed information processing.

Inspired in the dynamics of harvester ants, (Gordon *et al.* 1992) proposed a neural network model of task allocation where individual ants are represented by a sequence of Boolean variables instead of a single ON-OFF description. Observations from extensive field work on harvester ants (*Pogonomyrmex*) show that members of an ant colony perform a variety tasks outside the nest, such as foraging and nest maintenance work. Remarkably, this is a monomorphic species, i.e. individuals exhibit identical phenotypes. The number of ants actively performing each task changes over time due to task switching as well as the presence of inactive workers (Gordon 1986). As discussed in (Gordon 2010) interactions among ants involve physical contact. This allows sensing the state of other nestmates allows to create a network of information exchanges. Experimental perturbation of the number of ants performing a given task triggers changes in the numbers of individuals performing other tasks. Importantly, this switching dynamics is a consequence of the microscopic, local ant-ant interactions. The attractors associated to normal and perturbed conditions is thus a collective-level outcome of individual interactions.

In their model, Gordon and co-workers consider a set of four main tasks. This choice is partially due to the observation of four kinds of tasks, namely: patrollers, foragers, nest maintenance and midden workers displayed by harvester ants. Additionally, individuals can become inactive (as reported in ant colonies, see previous section). Since each type of ant performing any of the four tasks can become inactive, the model assumes that eight possible vectors can represent the available space state which can be covered by an internal state of three binary variables (Gordon *et al.* 1992). Specifically, ants are described now as 3-spin vectors 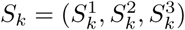. In their original paper, they use the notation *P* = active patroller, *F* = active forager, *N* = active nest maintenance worker and *M* = active midden worker. The lower case versions (*p, f, n, m*) would indicate inactive versions of the previous vectors. The space of possible internal states is indicated in Fig. 6a-b. These are represented as vertices of a Boolean cube, where all states are respectively indicated as strings of +1 and −1 values.

**Fig. 6:**
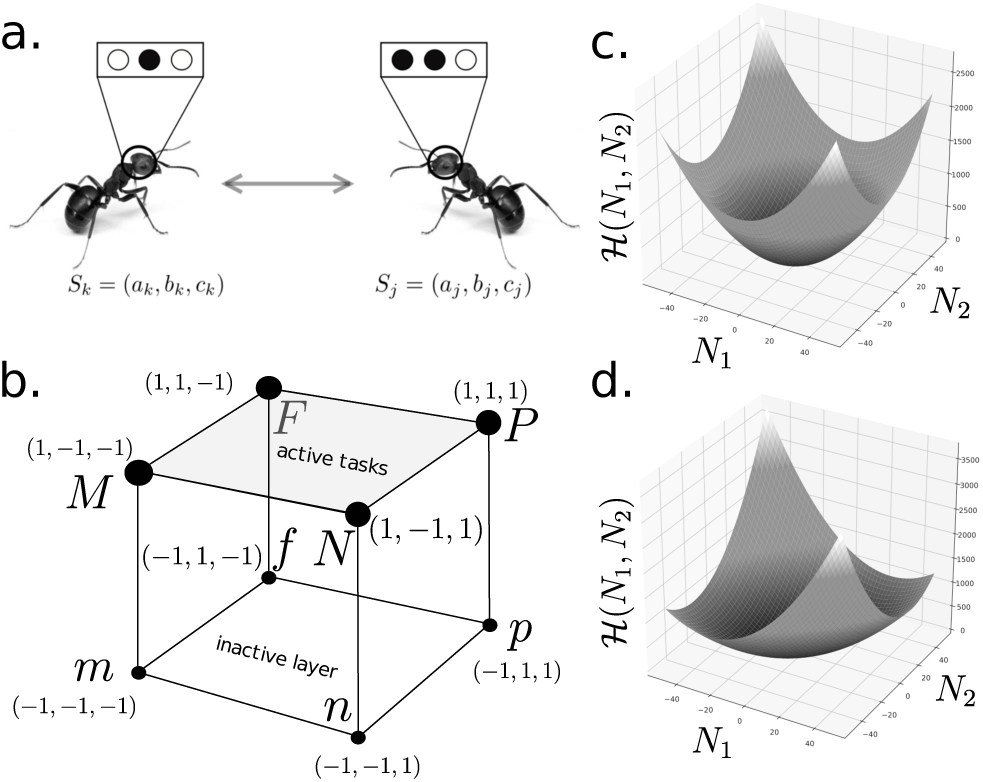
Neural network model of task allocation in ant colonies. The dynamics of harvester ants in (Gordon *et al.* 1992) can be described in terms of virtual ants (a) each carrying a 3-spin internal description, with changes taking place by means of direct pairwise interactions. The total state space is a three-dimensional Boolean cube (b) where we indicate active (observable) tasks in the top of the cube while a lower layer of inactive states is formed by a flip in the first spin (negative for inactive ants). The model exhibits an attractor dynamics with an associated potential (energy) function. Displayed in (c), the potential function is easily found for a two-task system for a specific (symmetric) values of parameters.

The simplest approach for this problem is to assume that the different components of the internal state act independently, with different associated weight matrices. In this way, we would have

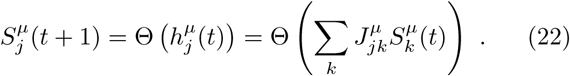

The (internal) state of 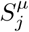 will remain stable after one interaction, provided that 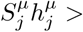 0. An energy function is defined accordingly as follows:

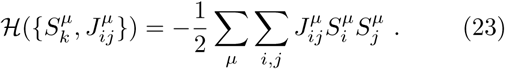

In the macroscopic realm, the observable state is the number of ants performing each task from the repertoire. It is then desirable to have a description where the energy minimisation is defined in terms of the set {*n_k_*}. Thus, the energy function now reads as

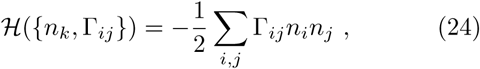

with a new set of parameters {Γ_*ij*_} that depend on the microscopic couplings and can be derived from the initial matrices (Gordon *et al.* 1992). This energy function allows a description of the system’s equilibrium states (attractors) as a high-dimensional surface which minima corresponds to the task allocation solutions. This form is consistent with a reaction-based dynamics where pairwise interactions among classes of individuals conditions the global dynamics. As a simple illustration of this idea, let us consider a two-state/two-task case, where it is not difficult to show that the energy function will correspond to

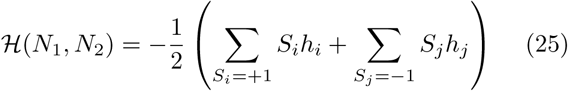

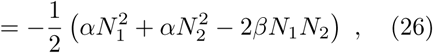

where we use Γ_11_ = Γ_22_ = *α* and Γ_12_ = Γ_21_ = *β*. We can easily recognise in this solution the elliptic paraboloid, displaying a single minimum. In Figure 6c we show an almost symmetric energy surface for *α* = 1, *β* = 0.1, whereas a less symmetric case is displayed in Figure 6d, where *β* = 0.5. In the latter, the coupling between the two tasks creates an elongated valley that would allow for more population fluctuations.

This model, with all the oversimplifications it contains, provides an elegant illustration of a major difference that might separate liquid from solid cognitive networks. The proper functionality of an ant colony, like the one described above, is satisfied for a given distribution of individuals performing the set of required tasks. Task allocation is thus achieved as a coarse-grained solution with high degeneracy: there are many ways to allocate individuals into given tasks. Thus the ant-ant interactions, although describable in terms of the standard threshold dynamics model, only provide a way to achieve the optimal state associated to the energy minimum in the task space.

#### 4. Collective dynamics of communicating populations

Insect colonies use different organic molecules (pheromones) to transmit signals and process information at a colony level. It is safe to assume that evolution the local environment for each individual ant, however, such an picture is incomplete when confronted to the full complexity of the colony. It is indeed the cobweb of diffusing pheromone-signals and ants acting as rewiring agents that confers the colony its true evolutionary potency. Individual ants are relegated to acting merely as cogwheels for the macrocospical system (Wilson 2012). This multiple-scale interrelation is the object of study of the present model by Mikhailov (Mikhailov 1993).

Suppose a colony of ants individually labeled as *i* = 1, …, N. Now, introduce two-state variables for each ant as *S*_*i*_ ∈ {−1, +1}, ∀*i*. Thus, vector 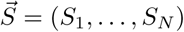 characterizes the full configuration of the system. In this model, ants are again acting as neural agents but they are also able to send out and receive messages into and from the colony. A message is encoded in a pheromone cocktail, and ants continuously secrete it. To simplify the system’s dynamics we will consider that a *message* is fully described with two labels, namely *µ*_*σ,j*_ = (*σ, j*), where *σ* ∈ {−1, +1} and *j* corresponds to the address tag. In other words, message *µ*_*σ,j*_ delivers information *σ* to the *j* − *th* ant.

During a time interval *τ*, multiple messages are sent all over the colony. Define *m*(*σ, i*) = ∑_*µ*|*τ*_*µ*_*σ, i*_, as the sum of all +/− signals tagging ant i over time interval *τ*, respectively (Fig. 7a). We then impose the following dynamics on the ant-states:

**Fig. 7:**
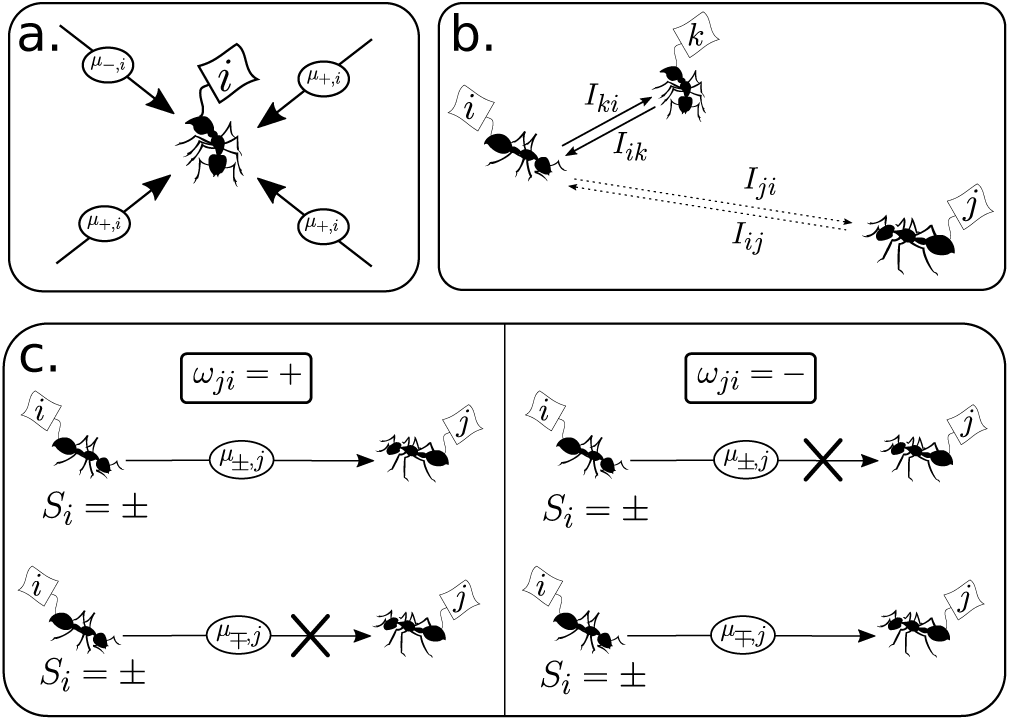
Collective communication dynamics in ant colonies. In (a) we display an agent *i* and a set of messages reaching it within time *τ*, all addressed to *i* while some carrying the + order others the order. These messages will be integrated according to (27). On the other hand, (b) shows how interactions via message sending depends on the frequency (or intensity) of messaging between agents, *I*. Notice that *I* values decay with the distance. Finally, the way that orders are sent by senders (c) depends on yet another set of couplings {*ω*_*ij*_ ∈ {−, +}}, which determine whether a + or a − order will be dumped into the system depending on the actual state of the sender *S*_*i*_ = ±. Schematically, the arrow connecting sender and receptor is blocked (crossed out) for anticorrelated correlation between coupling *ω*_*ji*_ and sender state *S*_*i*_.

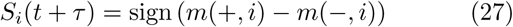

Let us introduce a *correspondence matrix*, {*ω*_ij_}, with each of the *N* (*N* − 1) elements of the former taking values {−, +}. The function of this matrix is to determine whether a signal will be sent or not in a time interval *τ*. The way it works is depicted in Figure 7c. If *ω*_*ji*_ = +, the sender ant, *i*, will send a message *µ*_±,*j*_ to ant *j* only if *S*_*i*_ = *±*, whereas for *ω*_*ji*_ = *−*, the message will be anti-correlated with the state of *i*, i.e., a message *µ*_±,*j*_ is sent only if *S*_*i*_ = ±. In simpler terms, the correspondence matrix distinguished two channels of information transfer: correlated (*ω* = +) or anticorrelated (*ω* =) message and sender-state. On the other hand, we define a *frequency distribution*, *I*_*ij*_, as the number of messages per unit time *τ* that ant *j* is sending to ant *i* (see Fig. 7b). Within a spatial context, it is clear that *I*_*ij*_ = *I*(|*i* − *j*|), where | *·* | is the distance between two ants.

Withall, let us consider the dynamics of the messages present in the system with labels (*σ, i*),

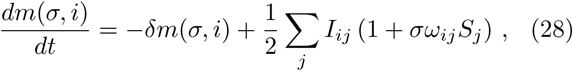

where we have dubbed *δ* the message decay rate. Therefore, at the stationary regime, we expect

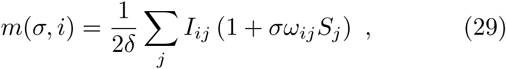

which, combined with (27), leads to

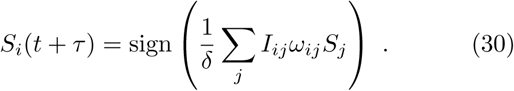

Notice that (30) is equivalent to the Hopfield model (1), provided that *ω*_*ij*_ = *ω*_*ji*_. Thus, patterns can be stored in a similar fashion by following a Hebbian approach by associating

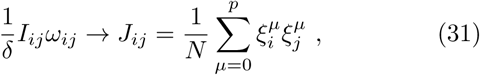

where, as in section II.A, 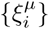 will correspond to the agent states of *µ* = 1, …, *p* different stored patterns. Although limitations to capacity will also apply here, perhaps more interestingly, other constrains will too arise, namely:

1. Agent-to-agent distance dependence on the signal intensity, *I*_*ij*_ = *I*(|*i − j*|), which should be take the form of a monotonically decreasing function. Effectively, this leads to a *diluted* network, i.e., every agent does not connect to every other agent.
2. Environmental noise: signal loss due to fluctuations of the information channel. This can be formalized as thermal noise, which has also been discussed in section II.A.
3. Cost-efficiency effects: the adress-message system devised here carries with it a large cost on the senders to produce the necessary chemical repertoire so that the signal is well-transmitted with minimal error. Below, we will discuss further on how to address these problems and figure out their implications in a collective computational levels.

### B. The Immune System as a Liquid Brain

The Immune System (IS) consists of a myriad of chemical compounds (e.g. antibodies, cytokines) and multiple cell lines (B-cells & T-cells or lymphocytes, macrophages, etc.) aggregated into a multi-component complex system. The essential purpose of the IS is to detect external and malicious agents (antigens) such as viruses, bacteria or cancerous cells; and prompt an according reaction (antigen neutralisation or tolerance). At the same time, it must be able to distinguish the latter from internal signals (the *self*). As such, the IS must be capable of processing, storing and manipulating large amounts of information (Delves & Roitt 2000).

The map of interactions of the IS can be depicted as an interwoven web of signalling and response functions between all its agents. Unravelling a full picture of the IS is beyond the scope of this work. For the purpose of our discussion, we will focus on the three core elements that significantly shape the IS architecture: T-Cells, B-Cells and Antibodies (Ab). Lymphocytes have specific enzymes on their membranes that store a molecular compound that has been randomly generated during its maturation process. This compound binds to some specific fragments of proteins (epitopes) coming from an antigen (often through an antigen presenting cell), hence prompting an internal cascade of reactions that activate the lymphocyte. The collection of receptors of a given lymphocyte clone-line is dubbed an *idiotype*.

Upon detection, B-Cells (aided by helper T-Cells) will proliferate thus generating copies of the same receptor structre, while secreting large concentrations of its specific antibody. In summary, the clonal expansion theory (Burnet 1959) states that, since the generated clones share their idiotype, successive binding to the antigen will be triggered and an amplification process will lead to immune response (Perelson & Weisbuch 1997).

On the other hand, a more systemic approach to the IS reveals an underlying network of idiotypes that excite or inhibit one-another through the *same* detection/reaction mechanisms as with antigens. This penomenon is known as an *idiotypic cascade*: an initial perturbation (antigen) activates a series of idiotypes filling the system with their corresponding antibodies (Ab1), which, in turn, are detected through by another set of idiotypes thus prompting a second batch of antibodies (Ab2) and so on and so forth. This observation suggests a network scheme where each node is associated with an idiotype and each link will correspond to an interaction between any two idiotypes (see Fig. 5a-c).

Idiotypic cascades were first observed and theorized by Jerne (Jerne 1974) and have since spurred a scientific debate between the *allopoietic/autopoietic* (reductionist/systemic) approaches to the IS (Barra & Agliari 2010, Perelson 1989, Parisi 1990). While Burnet’s theory provides some mechanisms for how the IS generates its idiotypic repertoire capable of self/non-self discrimination, Jerne’s network approach complements this process and shows how a distributed computation concatenated to clonal theory might give rise to crucial information-processing aspects of the immune response.

In this section we will study some fundamental aspects of the IS as a liquid brain. We will begin by looking at the size of the IS and how it is constrained by its fundamental function of antigen detection and discrimination. Then we will study how the IS is capabe of storing information at a network level, discuss how it makes use of its idiotypic landscape structure to naturally reproduce a reliable self/non-self classification and briefly comment on the implications of such a systems-view on the IS.

#### 1. Simple constrains for the probability of detection

Early studies of the IS showed that epitope reactivity for a generic lymphocyte (B-cell or T-cell) is of the order 10^*−*5^, in other words, the probability that a random epitope binds to the surface of a lymphocyte is given by *p* ≃ 10^*−*5^ (Perelson & Weisbuch 1997). This begs the question: why would not the IS organize such that *p* ~ 1?

In (Percus *et al.* 1993), a simple argument was put forward to show that the fact we observe such values of *p* might be related to the problem of self/non-self recognition, which strongly constrains the way the IS is assembled.

Consider the following definitions: *n* is the total number of receptors in the IS repertoire, *N* is the number of foreing-epitopes for a given environment and *N*′ denotes the number of self-epitopes, or epitopes derived from cells belonging to the organism. Thus, the goal of the IS is to properly distinguish the foreing-epitopes while avoiding an immune response for the self-originated ones. Let us denote by *P* (*N, N*′ *n*) the probability that the repertoire of size *n* is able to properly detect *N* foreing-epitopes and not detect *N*′ self-epitopes. Note that the probability of non-recognition of a random epitope for a single lymphocyte is given by *q* = 1 − *p*. Hence,

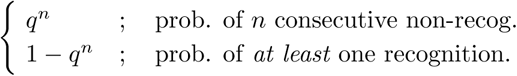

Therefore, we may now compute

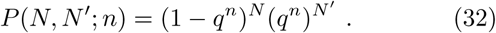

The goal is to maximize (32). This is easily done by maximizing the log *P* (*N, N*′; *n*), which leads to an optimal value for *q*

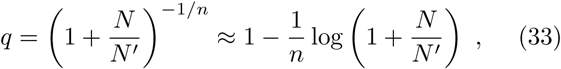

where we expanded the previous expression using 1/*n* ≪ 1. Notice that we can now write

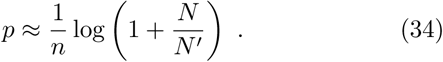

Early estimations of the repertoire size of a healthy human IS found *n* ~ 10^6^, while *N/N*′ ~ 10^10^ (see Perelson & Weisbuch 1997 pp. 1225-1229 and references therein). Along with *p* ∼ 10^*−*5^, such empirical values are compatible with the bounds imposed by (34). On the other hand, expanding (34) shows that the dependence on the ratio between self/non-self epitopes turns out to be very coarse, as *p* ~ (1/*n*) *N/N*′ + *O* (*N/N*′)^2^. More constrains on the complexity of the epitope molecule chains and the surface receptors requiere other sophisticated approaches (see Percus *et al.* 1992). In summary, the study of matching probabilities in detector systems such as the IS provides a robust understanding of the possible assemblies and architectures for such biological structures.

### 2. Percolation thresholds in the IS

After Jerne’s discovery of idiotypic cascades, novel ideas were put forward in trying to understand the organisational principles of the IS as a network. Perelson (Perelson 1989) introduced a simple model of the idiotypic cascading phenomenon. Given a repertoire of *n* idiotypes, i.e., *n* different types of antibodies, and assuming that paratopes and epitopes can be thought of as bit-strings of size *L* (see Fig. 8c), then we will consider that an antibody can detect (bind to) a given string if the number of matched pairs of the ordered paratope-epitope interaction exceeds a threshold value, *θ < n*. As we will see, this readily imposes a strong bounds on the system performance.

**Fig. 8:**
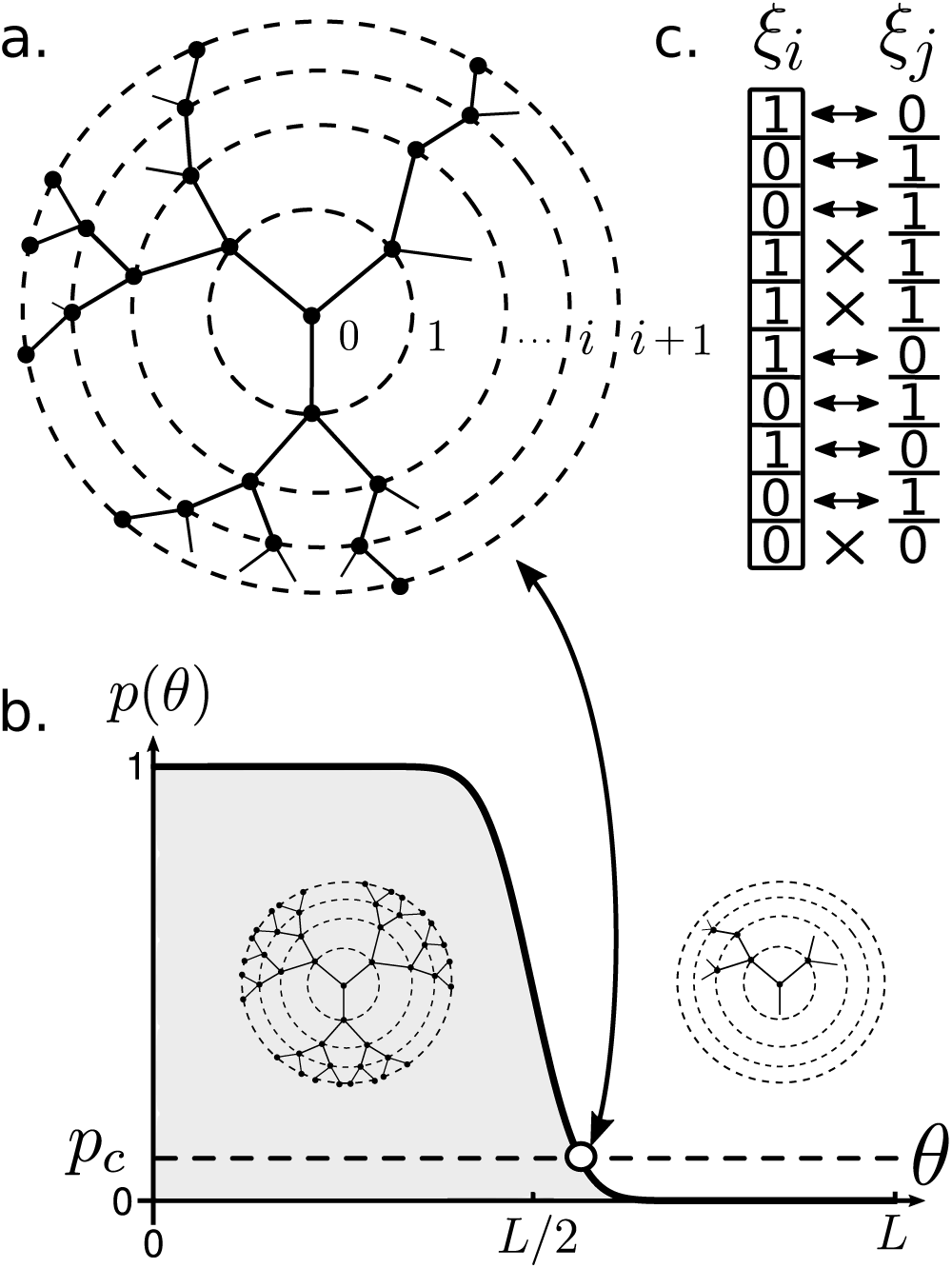
Percolation in immune networks. Idiotypic cascades take place at a network level in the IS. In (a) a critical percolation cascading on a Bethe lattice of degree *z* = 3 is shown. Concentric circles delimit successive layers of the cascade. The percolation probability depends on the the matching threshold *θ* as shown in (b). At low threshold values the system is highly connected, allowing deep penetration across layers, while for high *θ*, the matching probability decays abruptly, leading to a phase of low connectivity with small-sized cascades. Right in the interface, we have the percolation point. Finally (c) portrays two strings (eptiopeparatope) of length *L* = 10 with 7 matching pairs and 3 non-matching pairs. For example, if threshold *θ* = 5, this particular pair of strings would react, whereas for high fidelity matching (*θ* = 8), the pair would not connect.

Recall that, under the Jerne’s paradigm, antibodies are now capable of matching with other antibody types and concatenate into an idiotypic cascade. Thus, we can infer that, for a high threshold value (low reactivity), less antibodies will be matching, but also less antibodies will be able to detect and react to a given antigen. On the other hand, the reverse is also true: for low values of *θ* (high reactivity), antibodies will be triggered altogether, as the matching probability is expected to increase. Therefore, it is interesting to study what type structure will emerge from this simplified model.

Suppose that a given antibody is physically connected to a number of antibodies *z*, i.e., it will encounter up to *z* other antibody types but might or might not bind to them. Now, the probability that any pair of antibodies *do* match is denoted by *p*, which, by definition, will depend on *θ* (see below). Thus, given an initial perturbation into the system (such as antigen exposure) then an idiotypic cascade is triggered, where idiotypes react to eachother. Such a process will look like a Bethe lattice of degree *z* (see Fig. 8a). Denote by 𝓐(*i*) the number of activated antibodies at the *i*–th layer of the tree, then it is easy to show that:

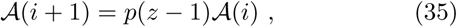

which implies that there will by a characteristical probability value *p* = *p*_*c*_ = (*z* − 1)^−1^, at which the network becomes connected exhibiting a *percolation* phase transition (Solé 2011). For values of *p* > *p*_*c*_, the network is fully connected, while for *p < p*_*c*_, any initial perturbation will eventually die out (see Fig. 8a-b).

For the IS one can argue that *z ~ n*, in other words, the system is sufficiently fluid and the coarse number of elements is sufficiently large so that any physical interaction can occur. This sets a value on the critical threshold at *p*_*c*_ ~ n^−1^. On the other hand, one can compute *p* = *p*(*θ*) by assuming that each bit, out of the *L*-sized strings, is generated by a coin toss. Then, the probability of having two strings with sufficient complementary bit-to-bit values is

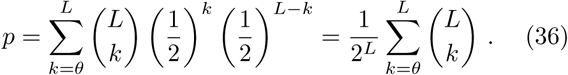

which is plotted in Fig. 8b. We observe a sudden transition from low to high reactivity at around *θ ~ L*/2. In fact, as *L* → ∞, then *p*(*θ*) → 1 − Θ(*L*/2).

Both *n* and *p* have been independently measured (Perelson 1989 pp. 19-20 and references therein). The repertoire size is estimated to be of the order *n* ~ 10^6^, while *p* ~ 10^−5^. Hence, the IS operates in the post-critical regime, where connectivity is high and large cascading events are common.

#### 3. Information storage in the immune networks

In the search for a clear understanding of how the IS stores and process information, optimisation arguments as above do not suffice under the light of Jerne’s theory of idiotypic networks. Initial attempts to describe how information is distributed over the network connecting different idiotypes were put forward by De Boer, Hogeweg, Weisbuch and Perelson (see Wiesbuch & Perelson 1997, pp. 1229-1258 and references therein). Here, we will briefly summarize a minimal model by Parisi (Parisi 1990) that involves Hopfield-like NN and imposes global limits on the pattern recognition processes that a distributed network of idiotypes must follow.

Consider the set of antibody binary concentrations {*c*_*i*_(*t*) ∈ {0, 1}}, for *i* = 1*, …, N*, with *N* the total number of antibodies of a healthy human IS (around 10^6^ − 10^7^). To all effects and purposes, *antibodies* and *idiotypes* are interchangable from here onwards. Next, we model idiotypic interaction networks, by imposing a dynamical process of idiotype concentrations in the same spirit of (1):

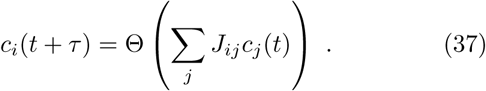

Now, the interactions between different idiotypes are mediated by {*J*_*ij*_}, for which we consider the following properties:

a. *J*_*ij*_ = 0, i.e., no idiotype self-interaction is allowed, which is the case for paratope-epitope complementarity matching.
b. *J*_*ij*_ = *J*_*ji*_, which is a simplification of the Onsager affinity relations between idiotypes [101], log |*J*_*ij*_| = log |*J*_*ji*_|.
c. *J*_*ij*_ = *U* (−1, +1), *∀i* ≠ *j*.

Condition (*c*) states that the values of the off-diagonal elements of *J*_*ij*_ are taken from the uniform distribution between [−1, 1]. These approximation allows for a derivation of overall limits of distributed storage of information. The system is now described as a spin glass (Amit *et al.* 1985, Sompolinsky 1988, Mezard *et al.* 1987).

Stable solutions for this particular problem turn out to be fully characterized by an average number of *pre-assigned concentrations*, *M*. In other words, a generic initial configuration of concentrations will inevitably *flow* into a stable state by switching concentration values on and off until a pre-assigned configuration of concentration levels is reached. These global stable states act as memory basins similarly to how memory is stored in the aforementioned NN models. Naturally, *M < N*, thus, we can define *α* < 1 such that *M* = *αN*.

Spin glass theory (Mézard *et al.* 1987) predicts that, for *N* ≫ 1, out of the total 2^*N*^ possible binary states of the system, and for conditions (*a*) − (*c*), a total of 2^*λN*^ patterns can be stored, with *λ* ~ 0.3. Withall, we can now try to understand the relation between *λ* and parameter *α*.

Let us consider the probability (*p*_*m*_) of randomly choosing a “memorized state” out of all the possible configurations or, simply, *p*_*m*_ = 2^*λN*^/2^*N*^ = 2^−(1−λ)*N*^. However, because only *M* pre-assinged concentrations are required to fully describe an attractor, we then expect a number of compatible solutions per stable state. Thus, let us compute the average number of compatible solutions per attractor as

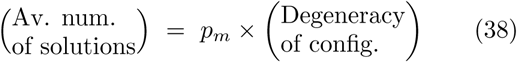

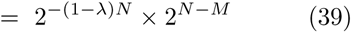

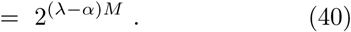

Notice that the average number of solutions will be greater or equal to one iff *α < λ* ~ 0.3. Essentially, this imposes a bound in *M*. In other words, if we denote *α*_*c*_ = *λ*, then for *M > α*_*c*_*N*, no equilibrium states are found. Thus, *M*_*c*_ ≡ *α*_*c*_*N* is the maximum number of pre-assigned antibody concentrations such that the dynamics imposed by (37) flow into well-defined stored patterns. This effectively constrains the memory content that an idiotypic interaction web is able to store.

A major insight from this model by Parisi is the fact that selective preassure goes beyond the genetic component involved in epitope/idiotope generation. Indeed, the reductionist approach is insufficient in trying to capture the full picture of IS evolution, as the information processing and storage occurring at the idiotypic network scale involves a higher order level at which selection will too operate.

#### 4. Idiotypic networks as liquid neural nets

In the remaining of this section we will outline a model by Barra & Agliari (BA) (Barra & Agliari 2010) based on statistical physics of a well-mixed/liquid neural web representing Jerne’s idiotypic network. Let us assume:

i. A given clone idiotype is fully characterised by a string of *L* bits. All idiotypes are of the same size.
ii. Each string is obtained from successive, independent coin-tosses with values {0, 1}.
iii. The number of cells of a clone-type is sufficiently large so that potential idiotypic interactions are always carried out with their respective intensity values.

Assumptions (*i*) − (*ii*) are sensible first approximations to the biological processes the IS undergoes during maturation (Delves & Roitt 2000). On the other hand, a sufficienty high number of lymphocytes per idiotype is not realistic under the lights of clonal expansion theory. However, the goal of the BA model is to figure out the overall implications of having an idiotypic network description[102].

Let us construct an idiotope space Υ_*L*_ ≡ {ξ ∈ {0, 1}^L^} spanning all possible strings with bit-size *L*. Indexes *i, j*, … ∈ {1, …, N}, with *N* corresponding to the total number of different clone-types in the IS. A priori, a complete repertoire would seem to scale as *N* ~ 2^*L*^, however, as we will see, the network constrains will give rise to another scaling behaviour between the repertoire size and epitope/paratope length.

Next, we construct the network following a simple model of chemical complementarity. As usual, let us define a *complementarity function*:

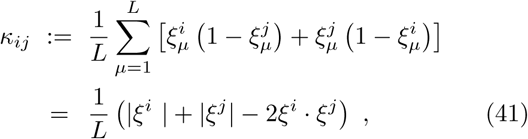

with 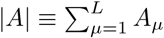. This accounts for the total number of complementary inputs between idiotypes *i* and *j*. E.g., suppose *L* = 5, then for *ξ*^*i*^ = (10101) and *ξ*^*j*^ = (01011), 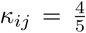 (Fig. 9b). In turn, this allows to construct a *chemical affinity function*

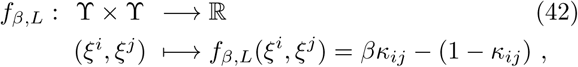

which is defined as a balance between repulsion and attraction effects of anti-complementary and complementary bit-pairs, moduled by trade-off parameter *β* ≥ 0. Thus, it will be bounded as −1 ≤ *f*_*β,L*_ ≤ + *β*, distinguishing two interactive regimes for each pair of idiotypes:

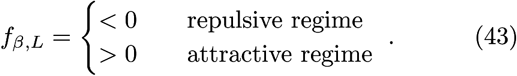

**Fig. 9:**
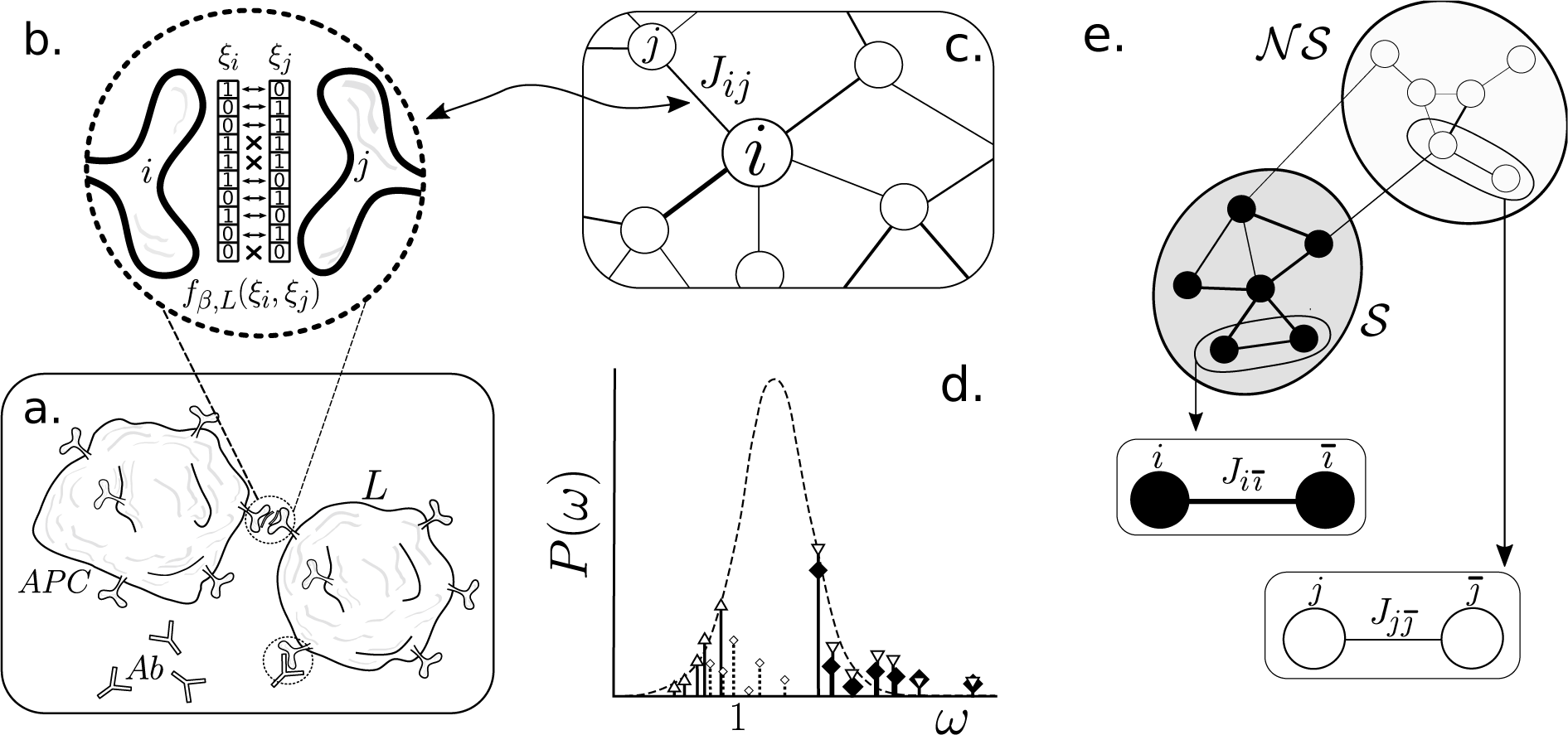
The IS as a liquid brain. (a) Represents an interaction between an antigen-presenting cell (APC), carrying a fragment of an antigen and presenting to a lymphocyte (L). Upon matching (b), the lymphocyte will react by secreting antibodies (Ab) with the corresponding matching code, thus flooding the system with its idiotypic information ang prompting an idiotypic cascade. Figure (c) is a representation of the subjacent idiotypic network operating accross the IS. This network is actually self-organised into two major blocks (d)-(e) of heavily influential (darker region) and weakly influential (lighter region) nodes. Such an effect can be computationally studied by looking at the strength distribution (d), *P* (*ω*), and noting that picking a random node from the right/left (strong/weak) (*i/j*) ends of the spectrum, and then looking at its corresponding next neighbor strength 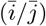, they typically fall under the same cathegory, i.e., strong/weak nodes connect to strong/weak nodes. This suggest an network-like mechanism for tackling the Self(𝓢)/Non-Self(𝓝𝓢) classification problem (*ω* axis is depicted in logarithmic scale). Strong nodes are responsible for self-adressed Ab, and viceversa. Figure (d) is adapted from (Barra & Agliari 2010).

Following these precepts, let us outline how the unweighted network of idiotype-idiotype interactions will unfold. The IS can be arguably approximated as a well mixed system. This means that, following (*iii*), any possible physical interaction (B-Cell/T-Cell or APC/T-Cell, etc.) occurs at a sufficiently high rate so that we need only to account for their internal affinity structure. Let us then define *p*_*β,L*_ as the probability that two generic idiotypes display a matching interaction. Consider the following:

1. The idiotype strings, {*ξ*^*i*^}, are extracted by a successive *L* random coin-tosses with equal probability for {0, 1} values, i.e. *p*_0_ = *p*_1_ = 1/2
2. The complementarity *κ*_*ij*_ and affinity *f*_*β,L*_ functions fully regulate the interactions. In particular, we define a link between two generic idiotypes (*ξ*^*i*^, *ξ*^*j*^) iff *f*_*β,L*_ (*ξ*^*i*^, *ξ*^*j*^) *>* 0, i.e., if the pair lays on the attractive regime.

In general, the probability for any two idiotypes to produce a complementarity value, *κ*, is 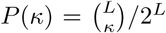 Now, owing to assumption 2 and (42), then

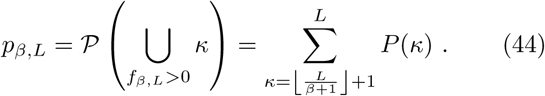

Since *N* is the total number of different idiotypes, the emergent network picture will be described by an Erdös-Renyi graph with degree distribution

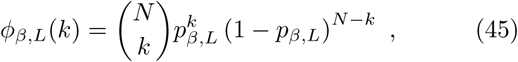

which mean value corresponds to 〈*k*〉 ≈ *p*_*β,L*_*N*. ER networks display a percolation point at which the system acquires a giant connected component (Solé 2011). Typically, this occurs at 〈*k*〉 = 1, associated to *p*_*c*_ := 1/*N*. Next, we explore what regime should we expect the idiotypic network to be in and how this reflects on the IS’s repertoire capacity.

For finite values of *L*, the shape of the function *p*_*β,L*_ as a function of *β* is that of a transfer function. Recall the trade-off parameter *β* separates the favourably repuslive regime (*β* < 1) from the favourably attractive one (*β* > 1), *β* = 1 corresponding to the symmetric case. Now, if chains (epitopes/idiotypes) are considered to be large, then an amplification process occurs depending on the favourably repulsive/attractive regimes determined by the value of *β*. Such amplification is reflected on the *switch*-like behaviour of the connection probability. On the other hand, since the percolation threshold will be of the order of 1/*N*, even if the system is repulsively-favoured (*β* < 1), it can still easily become fully connected.

Now, consider the three elements that are now comming together: probability of connection, *p*_*β,L*_; number of idiotypes (or different clones), *N*; and the average number of connections per idiotype, 〈*k*〉. While the probability of connection is purely a result from the internal chemical interactions, 〈*k*〉 is a defining feature of our network. Yet experimental data sets a value of *N* ~ 10^18^, and a connectivity between idiotypes in mature immune systems of *p*_*β,L*_ ~ 3 − 5% (Detours *et al.* 1996). This means that we should expect a densely connected network of around 〈*k*〉 ~ 10^12^.

Moreover, the fact that *p*_*β,L*_ is so low, suggests that the system operates at the repulsive regime. Withall, we may now compute the relation between epitope size *L* and number of idiotypes in the BA model by using (44) and *β* < 1. This results in a scaling relation 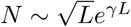, with *γ* < 1, as opposed to the bit-by-bit repertoire size, which would grow as 2^*L*^ (Barra & Agliari 2010).

Hitherto, we have been able to characterise idiotypic networks using only basic assumptions for chemical affinity, which has lead to a dense ER. But what kind of computations is this system able to perform? And how does the IS utilises its autopoietic features to distinguish the self/non-self? To provide an answer to these questions, we ought to look at a fine-grained version of the idiotypic network and inquire on how its interaction intensities are distributed over the net.

#### 5. Stewart-Varela-Coutinho theory

In their seminal papers Stewart-Varela-Coutinho (Stewart *et al.* 1989, Varela & Coutinho 1991) showed that a network systems approach to the idiotypic webs described by Jerne actually displayed two major interactive blocks: a highly connected (strong interacting) module and a loosely connected (low interactive) one. Such observation suggested that each module’s activity could correlate to the self and non-self reactions of the IS. More specifically, the strongly connected module acts as an auto-regulated dynamical subsystem that is continuously activated and auto-inhibited, this would correspond to a *tolerance* reponse, thus associated to the *self* (𝓢) stimuli, in a healthy IS. On the other hand, the low interactive module shows a basal activity in the system, but under the presence of a stimuli will be activated, thus prompting a *neutralisation* response. The latter module is then associated to the *non-self* (𝓝𝓢) stimuli. Hence, through this network structural property the vast repertoire of the IS is capable of sorting out the self/non-self.

Although this two-block structure could appear to be the result of an intricate evolutionary process, by following the BA model, a two-fold assembly akin to Stewart-Varela-Coutinho’s system is shown to emerge *for free*. This would suggest a generative mechanism capable of explaining the underlying Self/Non-Self modular structure independently of adaptive drives. Let us briefly explore how this phenomenon takes place at the weighted network level.

#### 6. Weighted idiotypic networks and mirror-types

The affinity function *f*_*β,L*_ works as a representation of the chimical reactions that take place on the cell surfaces, then, following a simple extension to the concept of interaction connection matrix *J*_*ij*_(*β, L*) can be defined as

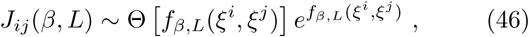

Notice how we still impose a lower threshhold of connectance by setting *J*_*ij*_ = 0 for *f*_*β,L*_ ≤ 0, which keeps the previous network picture, while turning on the matrix values smoothly on the attractive regime. The exact values for all the *J*_*ij*_ will depend on each realisation of the stochastically generated network of idiotypes {*ξ*^*i*^}, thus, it will be necessary to normalise each interaction parameter over all the space of possible networks (see Barra & Agliari 2010).

Once the interaction-intensities are at place, one can look at the total strength for each idiotype (node) as

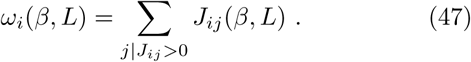

Heuristically, *J*_*ij*_ values characterize the robustness of a given *i − j* interaction, while *ω*_*i*_ measures how influential idiotype *i* is as relative to the whole network. Now, consider the weight frequency distribution *P*(*ω*), which can be shown to be well approximated by a normal distribution (Barra & Agliari 2010). Now, if we select an idiotype *i* and look at its first-order neighbors that inhibit *i*, namely *mirror-i* idiotypes (or simply 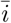), then it is possible to study how the system self-classifies these pairs into two major classes (see Fig. 6e): strong and weakly interacting pairs. The fact that the interaction is symmetrically strong/weak for each pair is a consequence of chemical complementarity in the affinity function.

However, this simple realisation turns out to be an extremely powerful tool precisely towards the self/nonself distinction. In summary:

- The *P* (*ω*) degree distribution separates the two regimes of strong/weak influential nodes (see Fig. 6d). The weak nodes (blank triangles) happen to have weak mirror-types (blank squares); whereas the strongly interacting nodes (reversed blank triangles) have mirror-types (black squares) that are too highly-interactive nodes. This mechanism gives rise to the *𝓢/𝓝,𝓢* modules.
- The strong block is hypothezised to account for the self-directed antibodies, while the weak module acts a basal signal only activated by the presence of non-self antibodies (triggered by external antigens).
- Establishment of robust memories occur more effectively at the weakly interacting block, as relative variations in the affinity values will produce more durable configuration changes in this network module.
- Autoimmune diseases can now be understood as deviations from the two-module structure, where strongly interacting circuits (responsible for self-addressed antibodies) may deviate towards lower weighted regions of the spectrum, thus triggering auto-immune response.

Hence, a natural mechanism for fundamental computational questions such as the Self/Non-Self identification is derived from first principles. This principles are constructed under the assumption that the interaction time-scales are small compared to the global observational time-scales, while, on the other hand the ability of the “neural agents” (idiotypes) to rapidly propagate throughout the environment ultimately allows the characterisation of the idiotypic network as a biologically meaningful system. Thus, the IS appears to be a limit case scenario for “liquid brains”, where it is precisely the high levels of agent-mobility that give rise to its capacity to solve classification problems.

This realisation spurs novel questions: are there a size limitations for the immune systems and their performance in terms of its physical embodiment? What are the consequences of these constrains onto the Self/NonSelf distinguishability? Or, in general, can different “immune systems” exists accross multiple scales?

## IV. DISCUSSION

The emergence of cognition in our biosphere has been marked by several key events that allowed the evolution of special classes of cell phenotypes along with ways of wiring them together. Nerve cells and nerve nets pervade the revolution towards new life forms capable of dealing with non-genetic information in complex ways. But the basic ingredients for the emergence of complex forms of information processing have appeared multiple times at different scales and in different evolutionary contexts (Balušska & Levin 2016). Neural-like processing systems have evolved as specialized organs but also as communities of moving agents. In both cases, agent-agent interactions involve some sort of recognition, internal communication coding and stimuli thresholds that decide if changes are made. As shown in previous sections, simple models can capture relevant phenomena associated to both classes of systems.

These two classes of networks share emergence as a major feature. Memory, learning or decision making are grounded in a set of bottom-up phenomena where emergent properties arise from individual, microscopic interactions (Fig. 10). Collective phenomena and a physics approach becomes a natural common field where to extract universal features. These emergent traits can be attractor basins associated to memory states or efficient task allocation, but can also be phase transitions due to the presence of critical connectivities or even criticality itself, enabling rapid and efficient information transfer.

**Fig. 10:**
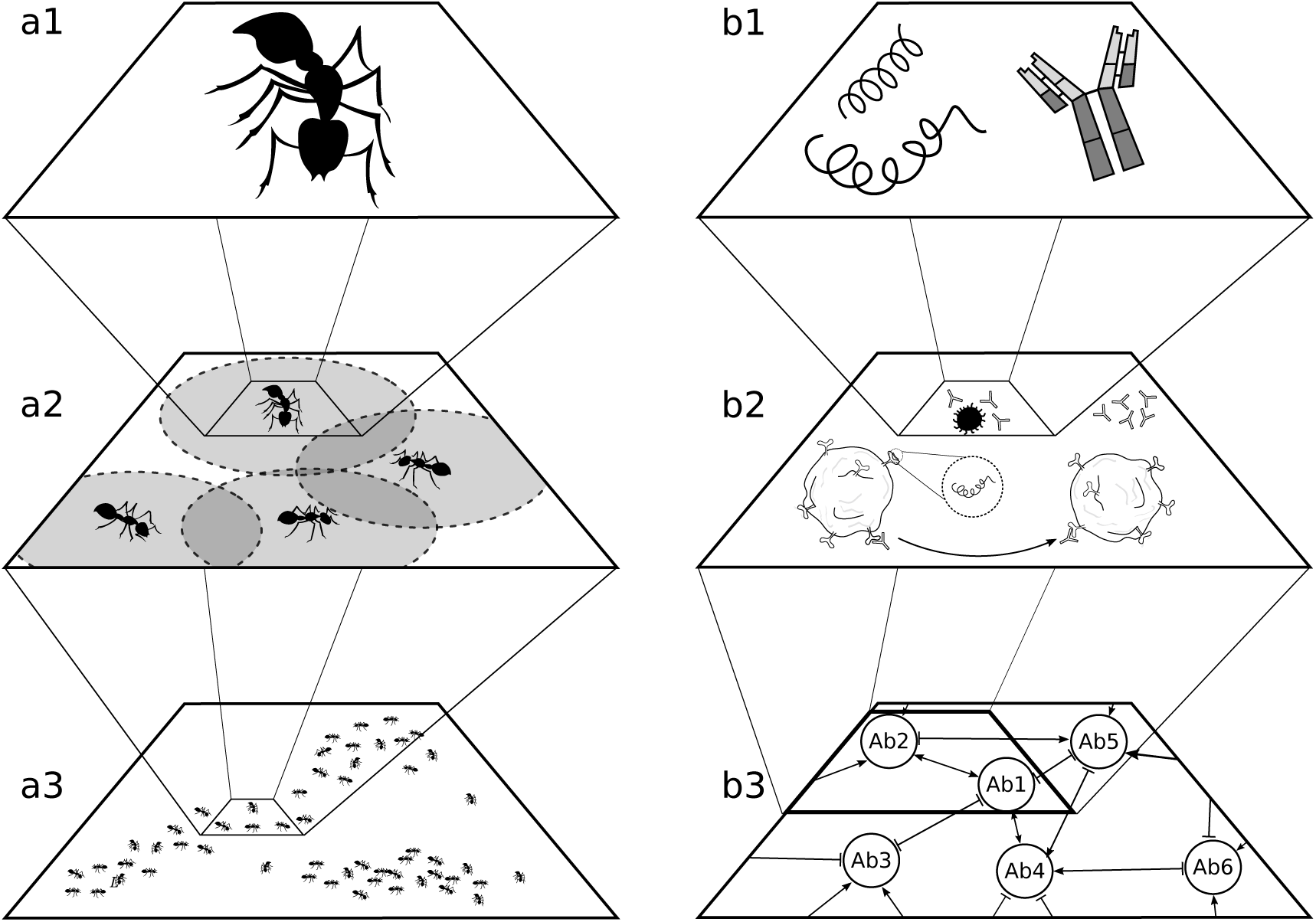
Multiscale dynamics in liquid brains. Each example of liquid brains involves, as it occurs with many other complex systems, several scales of description. Ant colonies (a) perform diverse functionalities, such as collective foraging (a3) on a colony-level basis. At a smaller scale, pairwise interactions among ants take place (a2). Such interactions are localised and, thus, constrained by spacio-temporal properties such as agent mobility or density. At the top of this hierarchy (a1) we encouter single ants as a system. These agents will be defined by a set of rules that drive their behaviour in this minimal scale. A similar scheme can be made for the immune system (b). Scales now involve a the idiotypic (or antibody type) network (b3), where information is processed, for instance, at the self/non-self discrimination level (see above). As we zoom in, we encounter the cellular-scale interactions level (b2), which are also associated to the simple-matching recognition dynamics. Finally, yet another level of complexity is reached at the description of the IS elementary agents (b1): viruses, paratopes, epitopes and surface repectors.

What is missing from our previous models? An important piece of complexity that has been ignored in this discussion is the internal complexity of the agents. This is not necessarily a limitation. When dealing with complex systems, we always need to cut the details up to some point in order to tackle the problem and provide true understanding. That is imposed by the kind of question being addressed. As with the Hopfield model and other classical neural network approaches, neuronal complexity is reduced to the minimal description. The immune system too is a rather sophisticated system, exhibiting a considerable diversity of cell types and interactions. Ants, on the other hand, are not simple strings of Boolean bits. But individual cognition exists: ants carry actual brains inside their heads, even if small ones (Webb 2012).

As pointed out by some authors (Dornhaus & Franks, 2008) even if these brains are orders of magnitude smaller than ours, they pervade some types of cognitive skills. How important are they compared to colony-level cognition? This is an open question that will require further attention. It might be the case that increases in the cognitive complexity of ant colonies is accompanied with reductions in individual’s cognition. Such trade-off has been explored in other contexts, like the evolution of multicellularity (McShea 2002), under the term *complexity drain*. Previous work use coupled discrete maps to suggest that collective computation might display too this phenomenon (Delgado & Solé, 1997b). Further multiscale models of liquid brains should be developed to properly address this question.

What other systems can be described as liquid brains? As mentioned before, computations arising from genegene regulatory links within cells have been studied using similar formal schemes, from Boolean tables to threshold networks (Kauffman 1984, Karlebach & Shamir 2008). Several observations also suggest that gene networks might be critical (Serra *et al.* 2007, Balleza *et al.* 2008, Torres-Sosa *et al.* 2009, Daniels *et al.* 2018).

A final comment needs to be made concerning with the physical phases used to present our classification between liquid and solid. The use of the term “liquid” to label the classes of systems discussed here is only partially appropriate. Particularly in relation with ants, their collective dynamics is more appropriately described as *active matter*: ants (as well as bacterial and robotic swarms) need to be understood as interacting self-propelled robots (Vicsek & Zafeiris 2012). Here too the statistical physics approach has splayed a key role in understanding coordinated behaviour and its transitions. Once again, in spite of considerable differences, deep analogies exist between classical equilibrium statistical physics systems and those made of active units. Understanding the cognitive complexity of liquid brains and its limits can provide deep insights into the evolution of information-processing, computational systems grounded in living structures.

## Ethics

N/A.

## Data Accessibility

This article has no additional data.

## Author’s Contributions

All authors built and analysed the mathematical models. All authors gave final approval for publication.

## Acknowledgments

The authors want to thank Luís Seoane, Sergi Valverde, Victor Maull, and the other members of the Complex Systems Lab for stimulating discussions. Special thanks to Bob Merriman for his inspiring ideas. We also thank the Santa Fe Institute for hosting the Working Group on “Liquid Brains, Solid Brains”.

## References

[1] Adamatzky A. Physarum machines: computers from slime mould. World Scientific, Singapore, Singapore (2010).

[2] Altshuler, E., Ramos, O., Núnez, Y., et al, 2005. Symmetry breaking in escaping ants. Am. Nat., 166, 643–649.

[3] Amé, J.M., Halloy, J., Rivault, C., Detrain, C. and Deneubourg, J.L., 2006. Collegial decision making based on social amplification leads to optimal group formation. Prod. Natl. Acad. Sci. USA 103, 5835–5840.

[4] Amit D.J., Gutfreund H., Sompolinsky H. 1985. Spinglass models of neural networks. Phys. Rev. A 32(2), 1007–1018.

[5] Balleza E., Alvarez-Buylla E.R., Chaos A., Kauffman S., Shmulevich I., Aldana M. 2008. Critical dynamics in genetic regulatory networks: examples from four kingdoms. PLoS One 3(6): e2456.

[6] Balušska F., Levin M. 2016. On having no head: cognition throughout biological systems. Front. Psychol. 7, 902.

[7] Barra A., Agliari E. 2010. A statistical mechanics approach to autopoietic immune networks. J. Stat. Mech. Theory Exp., P07004.

[8] Beckers R., Deneubourg J.L., Goss S. 1992. Trails and U-turns in the selection of a path by the ant Lasius niger. J. Theor. Biol. 159, 397–415.

[9] Benenson Y. 2012. Biomolecular computing systems: princples, progress and potential. Nat. Rev. Genet. 13, 455–468.

[10] Bonabeau E., Meyer C. 2001. Swarm intelligence a whole new way to think about business. Hardvard Business Rev. R0105G.

[11] Bonabeau E., Dorigo M., Theraulaz, G. Swarm intelligence: from natural to artificial systems. Oxford U. Press, New York, NY, USA (1999).

[12] Bonabeau E., Theraulaz G., Deneubourg J.L. 1998. The synchronization of recruitment-based activities in ants. BioSystems 45, 195–211.

[13] Bornholdt S. 2008. Boolean network models of cellular regulation: prospects and limitations. J. R. Soc. Interface 5, S85–S94.

[14] Bornholdt S. 2005. Less is more in modelling large genetic networks. Science 310, 449–451.

[15] Bray D. 1990. Intracellular signalling as a parallel distributed process. J. Theor. Biol. 143, 215–231.

[16] Brede M., Behn U. 2001. Architecture of idiotypic networks: percolation and scaling behavior. Phys. Rev. E 64, 011908.

[17] Burnet F.M. The clonal selection theory of acquired immunity. Vanderbilt U. Press, Nashville, TN, USA (1959).

[18] Buszaki G., Draguhn A. 2004. Neuronal oscillations in cortical networks. Science 304, 1926–1929.

[19] Chialvo D.R. 2004. Critial brain networks. Phys. A 340(4), 756–765.

[20] Chialvo D.R. 2010. Emergent complex neural dynamics. Nat. Phys. 6(10), 744–750.

[21] Cohen M.A., Grossberg S. 1983. Absolute stability of global pattern formation and parallel memory storage by competitive neural networks. IEEE Trans. Syst. Man. Cybern. 13(5), 815–826.

[22] Cole B.J. 1991. Short-term activity cycles in ants: generation of periodicity by worker interaction. Amer. Naturalist 137(2), 244–259.

[23] Daniels B.C., Kim H., Moore D., Zhou S., Smith H.B., Karas B., Kauffman S.A. & Walker S.I., 2018. Criticality Distinguishes the Ensemble of Biological Regulatory Networks. Phys. Rev. Lett. 12: 138102.

[24] Cole B.J., Cheshire D. 1996. Mobile cellular automata models of ant behavior: movement activity of Leptothorax allardycei. Amer. Naturalist 148(1), 1–15.

[25] Deco G., Jirsa V.K., Robinson P.A., Breakspear M., Friston K. 2008. The dynamic brain: from spiking neurons to neural masses and cortical fields. PLoS Comp. Biol. 4(8), e1000092.

[26] Delgado J., Solé R.V. 1997a. Noise induced transitions in fluid neural networks. Phys. Lett. A 229, 183–189.

[27] Delgado J., Solé R.V. 1997b. Collective-induced computation. Phys. Rev. E 55(3), 2338–2344.

[28] Delgado J., Solé R.V. 2000. Self-synchronization and task fulfilment in ant colonies. J. Theor. Biol. 205, 433–441.

[29] Delves P.J., Roitt I.M. 2000. The immune system. First of two parts. Adv. Immunol. 343(1), 37–49.

[30] Deneubourg J.L., Aron S., Goss S., Pasteels J.M. 1990. The self-organizing exploratory pattern ofthe argentine ant. J. Insect. Behav. 3(2), 159–168.

[31] Deneubourg J.L., Goss S. 1989. Collective patterns and decision-making. Ethol. Ecol. Evol. 1, 295–311.

[32] Detours V., Sulzer B., Perelson A. 1996. Size and connectivity of the idiotypic network are independent of the discreteness of the affinity distribution. J. Theor. Biol. 183, 409–416.

[33] Detrain C., Deneubourg J.L. 2006. Self-organized structures in a superorganism: do ants “behave” like molecules? Phys. Life Rev. 3, 162–187.

[34] Dorigo M., Birattari M., Stützle T. 2006. Ant colony optimization. IEEE Comp. Intell. Mag. 1, 28–39.

[35] Dornhaus, A. and Franks, N.R., 2008. Individual and collective cognition in ants and other insects (Hymenoptera: Formicidae). Myrmecol. News, 11, 215–226.

[36] Duarte A., Weissing F.J., Pen I., Keller L. 2011. An evolutionary perspective on self-organized division of labor in social insects. Annu. Rev. Ecol. Evol. Syst. 42, 91–110.

[37] Eckmann J.P., Feinerman O., Gruendlinger L., Moses E., Soriano J., Tlusty T. 2007. The physics of living neural networks. Phys. Rep. 449(1-3), 54–76.

[38] Forrest S. 1990. Emergent computation: self-organizing, collective, and cooperative phenomena in natural and artificial computing networks. Phys. D 42, 1–11.

[39] Forrest S., Hofmeyr S.A., Somayaji A. 1997. Computational immunology. Comm. of the ACM 40(10), 88–96.

[40] Farmer J.D. 1990. A rosetta stone for connectionism. Physica D 42, 153–187.

[41] Garnier S., Gautrais J., Theraulaz G. 2007. The biological princples of swarm intelligence. Swarm Intell. 1, 3–31.

[42] Goldenfeld N. Lectures on phase transitions and the renormalization group. Westview Press, Cambridge, MA, USA (1992).

[43] Gordon D.M. 1986. The dynamics of the daily round of the harvester ant colony (Pogonomyrmex barbatus). Anim. Behav. 34, 1402–1419.

[44] Gordon D.M. Ants at work: how insect society is organized. W.W. Norton & Co., New York, NY, USA (1999).

[45] Gordon D.M. Ant encounters. Princeton U. Press, Princeton, NJ, USA (2010).

[46] Gordon D.M., Goodwin B.C., Trainor L.E.H. 1992. A parallel distributed model of the behaviour of ant colonies. J. Theor. Biol. 156, 293–307.

[47] Goss S., Deneubourg J.L. 1988. Autocatalysis as a source of synchronised rythmical activity in social insects. Insectes Sociaux 35(3), 310–315.

[48] Hertz J., Krogh A., Palmer R.G. Introduction to the theory of neural computation. Vol I. Addison-Wesley, Redwood City, CA, USA (1991).

[49] Hesse J., Gross T. 2014. Self-organized criticality as a fundamental property of neural systems. Front. Neurosci. 8, 166.

[50] Wilson E.O., Hölldobler B. 2005. Eusociality: origin and consequences. Prod. Natl. Acad. Sci. USA 102(38), 13367–13371.

[51] Haken H. Synergetic computers and cognition: a topdown approach to neural nets. Springer-Verlag, Berlin, Germany (1991).

[52] Hölldobler B., Wilson E.O. The ants. Harvard U. Press, Cambridge, MA, USA (1990).

[53] Hopfield J.J. 1982. Neural networks and physical systems with emergent collective computational abilities. Proc. Natl. Acad. Sci. 79, 2554–2558.

[54] Hopfield J.J. 1994. Physics, computation and why biology looks so different. J. Theor. Biol. 171(1), 53–60.

[55] Jablonka E., Lamb M.J. 2005. The evolution of information in the major transitions. J. Theor. Biol. 239, 236–246.

[56] Jerne N.K. 1974. Towards a network theory of the immune system. Ann. Inst. Pasteur Immunol. 125C: 373–389.

[57] Karlebach G., Shamir R. 2008. Modelling and analysis of gene regulatory networks. Nat. Rev. Mol. Cell. Biol. 9(10), 770–780.

[58] Kauffman, S.A. The origins of order: self-organization and selection in evolution. Oxford U. Press, New York, NY, USA (1993).

[59] Kauffman, S.A., 1984. Emergent properties in random complex automata. Physica D 10, 145–156.

[60] McShea D.W. 2002. A complexity drain on cells in the evolution of multicellularity. Evolution 56(3), 441–452.

[61] McCulloch W.S., Pitts W. 1943. A logical calculus of the ideas immanent in nervous activity. Bull. Math. Biophys. 5, 115–133.

[62] Mezard M., Parisi G., Virasoro M.A. Spin glass theory and beyond: an introduction to the replica method and its applications. World Scientific, Singapore, Singapore (1987).

[63] Mikhailov A.S. 1993. Collective dynamics in models of communicating populations. In Interdisciplinary Approaches to Nonlinear Complex Systems, pp. 89–108. Springer Berlin Heidelberg.

[64] Millonas M.M., 1993. Swarms, phase transition, and collective intelligence; and a nonequilibrium statistical field theory of swarms and other spatially extended complex systems. SFI Working Papers 93–06-039.

[65] Miramontes O., 1995. Order-disorder transitions in the behavior of ant societies. Complexity 1(3), 56–60.

[66] Miramontes O., DeSouza O. 1996. The nonlinear dynamics of survival and social facilitation in termites. J. of Theor. Biol. 181, 373–380.

[67] Miramontes O., Solé R.V., Goodwin B.C. 1993. Collective behaviour of random-activated mobile cellular automata. Physica D 63, 145–160.

[68] Mora T., Bialek W. 2011. Are biological systems posied at criticality? J.Stat. Mech. 144, 268–302.

[69] Muñoz M.A. 2018 Colloquium: criticality and dynamical scaling in living systems. Rev. Mod. Phys. 90, 031001.

[70] Murray J.D. Mathematical biology. Springer-Verlag, New York, NY, USA (1989), pp. 610–650.

[71] Nicolis S.C., Deneubourg J.L. 1999. Emergent patterns and food recruitment in ants: an analytical study. J. Theor. Biol. 198, 575–592.

[72] Oster G.F., Wilson E.O. Caste and ecology in the social insects. Princeton U. Press, Princeton, NJ, USA (1978).

[73] Pagän O. Strange survivors: how organisms attack and defend in the game of life. BenBella Books, Dallas, TX, USA (2018).

[74] Parisi G. 1990. A simple model for the immune network. Proc. Natl. Acad. Sci. 87, 429–433.

[75] Percus J.K., Percus O.E., Perelson A.S. 1993. Predicting the size of T-Cell receptor and antibody combining region from consideration of efficient self-nonself discrimination. Proc. Natl. Acad. Sci. USA 90, 1691–1695.

[76] Perelson A.S., Weisbuch G. 1997. Immunology for physicists. Rev. Mod. Phys. 69(4), 1219–1267.

[77] Peretto, P., 1992. An introduction to the modeling of neural networks. Cambridge University Press.

[78] Perelson A.S. 1989. Immune network theory. Immunol. Rev. 110, 5–36.

[79] Plenz D., Niebur E., Schuster H.G. Criticality in neural systems. Wiley-VCH, Weinheim, Germany (2014).

[80] Rashevksy N. Mathematical biophysics: physico-mathematical foundations of biology. (3rd Ed.) Dover P., Inc., New York, NY, USA (1960).

[81] Rose S. The future of the brain. Oxford U. Press, New York, NY, USA (2006).

[82] Rosenblatt F. 1958. The perceptron: a probabilistic model for information storage and organization in the brain. Psych. Rev. 65 (6), 386–408.

[83] Serra R., Villani M., Graudenzi A., Kauffman, S.A. 2007. Why a simple model of genetic regulatory networks describes the distribution of avalanches in gene expression data. J. Theor. Biol. 246, 449–460.

[84] Strogatz S.H. Nonlinear dynamcs and chaos. Westview Press, Cambridge, MA, USA (1994).

[85] Solé R.V. Phase transitions. Princeton U. Press, Princeton, NJ, USA (2011).

[86] Solé R.V., Miramontes O., Goodwin B.C. 1993. Oscillations and chaos in ant societies. J. Theor. Biol. 161, 343–357.

[87] Solé R.V., Miramontes O. 1995. Information at the edge of chaos in fluid neural networks. Physica D 80, 171–180.

[88] Solé R., Amor D.R., Duran-Nebreda S., Conde-Pueyo N., Carbonell-Ballestero M., Montañez. 2016. Synthetic collective intelligence. BioSys 148, 47–61.

[89] Sompolinsky H. 1988. Statistical mechanics of neural networks. Phys. Today 41(12), 70–80.

[90] Stewart J., Varela F.J., Coutinho A. 1989. The relationship between connectivity and tolerance as revealed by computer simulation of the immune network: some lessons for an understanding of autoimmunity. J. of Autoimmunity 2(1), 15–23.

[91] Sumpter D.J.T. 2006. The principles of collective animal behaviour. Phil. Trans. R. Soc. B 361, 5–22.

[92] Tero A., Kobayashi R., Nakagaki T. 2007. A mathematical model for adaptive transport network in path finding by true slime mold. J. Theor. Biol. 244, 553–564.

[93] Torres-Sosa C., Huang S., Aldana M. 2012. Criticality is an emergent property of genetic networks that exhibit evolvability. PLoS Comp. Biol. 8: 1002669.

[94] Varela F.J, Coutinho A. 1991. Second generation immune networks. Immunol. Today 12(5), 159–166.

[95] Vicsek T., Zafeiris A. 2012. Collective motion. Phys. Rep. 517, 71–140.

[96] Webb B. 2012. Cognition in insects. Phil. Trans. R. Soc. B 367, 2715–2722.

[97] Weisbuch G. 1990. A shape space approach to the dynamics of the immune system. J. Theor. Biol. 143, 507–522.

[98] Wilson E.O. The social conquest of earth. W.W. Norton & Co., New York, NY, USA (2012).

[99] The same results are obtained when the active phase is used, since the two points just exchange their stability.

[100] A trade-off between polymorphism and pheromone repertory is evidenced, as caste differentiation already segregates behavioral states in a decisive way.

[101] Considering antysymmetric interactions leads to chaotic behaviour for the time-dependent dynamics, which is arguably not a good description of the IS as Ab concentrations would be observed to behave randomly.

[102] Assumption (iii) acquires more relevance in the full BA model as the coupling between the actual lymphocyte activity and the subjacent idiotypic network is studied. In this paper, though, we will only concern on the network-like features.

